# Thalamic involvement defines distinct slow-wave subtypes in NREM sleep

**DOI:** 10.1101/2025.01.16.633402

**Authors:** Damiana Bergamo, Giacomo Handjaras, Dante Picchioni, Emiliano Ricciardi, Guillaume Legendre, Pınar S. Özbay, Jacco A. de Zwart, Jeff H. Duyn, Giulio Bernardi, Monica Betta

**Affiliations:** IMT School for Advanced Studies Lucca, MoMiLab Research Unit, Lucca, Italy; University of Padova, Department of General Psychology, Padova, Italy; National Institutes of Health, National Institute of Neurological Disorders and Stroke, Advanced Magnetic Resonance Imaging Section, Bethesda, USA; University of Geneva, Department of Neuroscience, Geneva, France; Institute of Biomedical Engineering, Boğaziçi University, Istanbul, Türkiye

**Author notes:** **Correspondence:** Monica Betta, PhD - Giulio Bernardi, MD, PhD - IMT School for Advanced Studies Lucca MoMiLab Research Unit Piazza San Francesco, 19 Lucca - 55100, Italy. Equal last-author contribution (Jeff H. Duyn, Giulio Bernardi, Monica Betta).

**Keywords:** sleep, fMRI, EEG, thalamus, NREM, slow wave, arousal, autonomic

## Abstract

Slow waves (0.5–4 Hz) are a key feature of non-rapid-eye-movement (NREM) sleep, traditionally believed to arise from neocortical circuits. However, growing evidence suggests that subcortical structures, particularly the thalamus, may play a crucial role in initiating and synchronizing slow waves. We tested the hypothesis that single slow waves may arise from distinct cortico-cortical and thalamo-cortical mechanisms using simultaneous EEG-fMRI in healthy adults. Spatial mapping based on thalamic fMRI responses revealed two types of slow waves. The first type (C1) characterized by an early thalamic fMRI-signal increase, corresponded to large, efficiently synchronized waves associated with sleep spindles and with markers of higher arousal and autonomic activation. The second type (C2) is marked by an initial negative fMRI response and corresponds to smaller slow waves potentially resulting from cortico-cortical synchronization. These waves occur more often during phases of stable NREM sleep. These findings highlight distinct slow-wave subtypes with different thalamic involvement and, potentially, synchronization mechanisms.

## Introduction

The transition from wakefulness to non-rapid-eye-movement (NREM) sleep is marked by a shift from high-frequency, low-amplitude electroencephalographic (EEG) activity, to a pattern characterized by low-frequency, high-amplitude slow waves (0.5-4 Hz, ^1^). Sleep slow waves are thought to have a key role in the sensory disconnection and reduced level of consciousness characteristic of NREM sleep ^2–4^. Furthermore, growing evidence implicates NREM slow waves in sleep-dependent processes such as memory consolidation ^5^, synaptic strength homeostasis ^6^, and the clearance of metabolic waste products ^7–9^.

Traditionally, slow waves have been thought to originate primarily in neocortical circuits ^10,11^. However, recent findings from studies in animals ^12–16^ and humans ^17,18^ support a critical role of subcortical structures, especially the thalamus, in synchronizing and orchestrating cortical slow waves. A crucial unresolved question, though, concerns the possible existence of distinct slow-wave subtypes with unique features and generation mechanisms ^19–21^. One hypothesis posits the coexistence of slow-wave subtypes driven predominantly by cortico-cortical or subcortico-cortical synchronization ^3,22^. In this framework, diffusely projecting ascending systems may induce synchronized cortical neuronal off-periods, producing large, steep, and widespread slow waves. This mechanism aligns with the generation of K-complexes - large slow waves that can occur apparently spontaneously or in response to external stimuli ^23–26^. In contrast, small, shallow, and localized slow waves may arise from cortical generators, potentially reflecting less efficient cortico-cortical or cortico-thalamo-cortical synchronization. In the literature, a similar dissociation is made between slow oscillations—large, <1 Hz slow waves often associated with spindles—and delta waves, which are smaller and faster (1–4 Hz). These two types of slow waves exert distinct but complementary influences on memory consolidation^27^, suggesting that slow-wave classification is relevant not only mechanistically but also functionally ^28^.

In this study, we tested the hypothesis that slow waves are generated by distinct cortico-cortical and thalamo-cortical mechanisms. Using whole-night, simultaneous EEG-fMRI recordings in healthy adults, we employed thalamic blood-oxygen-level-dependent (BOLD) signal changes time-locked to slow wave onset as input for an unsupervised clustering analysis. This approach identified two distinct slow-wave sub-types (*clusters*) based on their associated thalamic activation patterns. We then systematically characterized these clusters in terms of morphology (amplitude, duration, and synchronization efficiency) and distribution across NREM sleep substages. Additionally, we examined the overlap between each cluster and classically defined K-complexes, as well as their association with sleep spindles (10–16 Hz) - waxing and waning oscillations commonly nested within the depolarizing phase of slow waves, indicative of thalamo-cortical interplay ^29–31^. We then explored the potential relationship between the two slow-wave clusters and peripheral vasoconstriction events linked to arousal-related changes in autonomic activity ^32–36^. Finally, we examined how these clusters relate to infraslow sigma oscillations, which are thought to reflect fluctuations in sleep stability and memory consolidation ^37,38^ . Our findings provide robust evidence for at least two slow-wave subtypes differing in their associated thalamic involvement. These subtypes exhibit distinct morphological characteristics, consistent with distinct generation mechanisms: one involving subcortico-cortical synchronization and the other primarily driven by cortico-cortical or cortico-thalamo-cortical synchronization. Furthermore, our analyses revealed the temporal dynamics underlying the generation of each wave type during NREM sleep. Thalamocortical waves predominantly occur during the descending phases of infra-slow oscillations—phases associated with higher arousal and increased sympathetic activity— whereas corticocortical waves generally emerge during the ascending phases, corresponding to more stable NREM sleep. This refined classification provides new evidence indicating that slow waves are not a homogeneous phenomenon and underscores the need for a more nuanced understanding of their regulation and functional roles. Notably, these subtypes may exhibit differential susceptibility to pathological alterations, and their characterization could provide novel insights into the contribution of sleep slow waves to neurological and psychiatric disorders.

## Results

We analyzed a total of 24 whole-night, simultaneous EEG-fMRI recordings obtained from 12 healthy adult participants (8 females, age range 18-34 years) acquired at the National Institutes of Health In Vivo NMR Research Center, USA (see ^39^). Each participant completed two experimental nights, yielding an average of 7.79 ± 1.41 EEG-fMRI acquired runs per night. Since our aim was not to assess homeostatic regulation across nights or first-night effect, but rather to maximize the amount of NREM sleep and the number of NREM graphoelements available for analysis, data from both nights were combined.

On average, each subject spent 464 ± 122 minutes (∼7.73 ± 2.04 hours) asleep, of which 426 ± 106 minutes in NREM sleep (N1= 49 ± 29 min; N2= 296 ± 84 min; N3= 81 ± 39 min) and 37 ± 23 minutes in REM sleep (Figure 1, Table 1; also see Supplementary Figure S1). We detected a total of 62,694 slow waves during NREM sleep, with a mean of 5,224 ± 2,135 slow waves per participant (Supplementary Figure S2, Table S1). Most slow waves (∼56%) were detected during N3 sleep, while ∼43% of them were identified in N2 sleep. We detected a total of 18,237 spindle events (1,519 ± 852 per subject), mostly during N2 sleep (∼75%; Supplementary Figure S3, Table S2). Finally, we identified a total of total 8,778 photopletysmographic pulse-wave amplitude (PWA)-drops reflecting peripheral vasoconstriction (731 ± 313 per subject), with about 12% events in N1, 72% in N2, and 16% in N3 (Supplementary Figure S4, Table S3).

**Figure 1.**
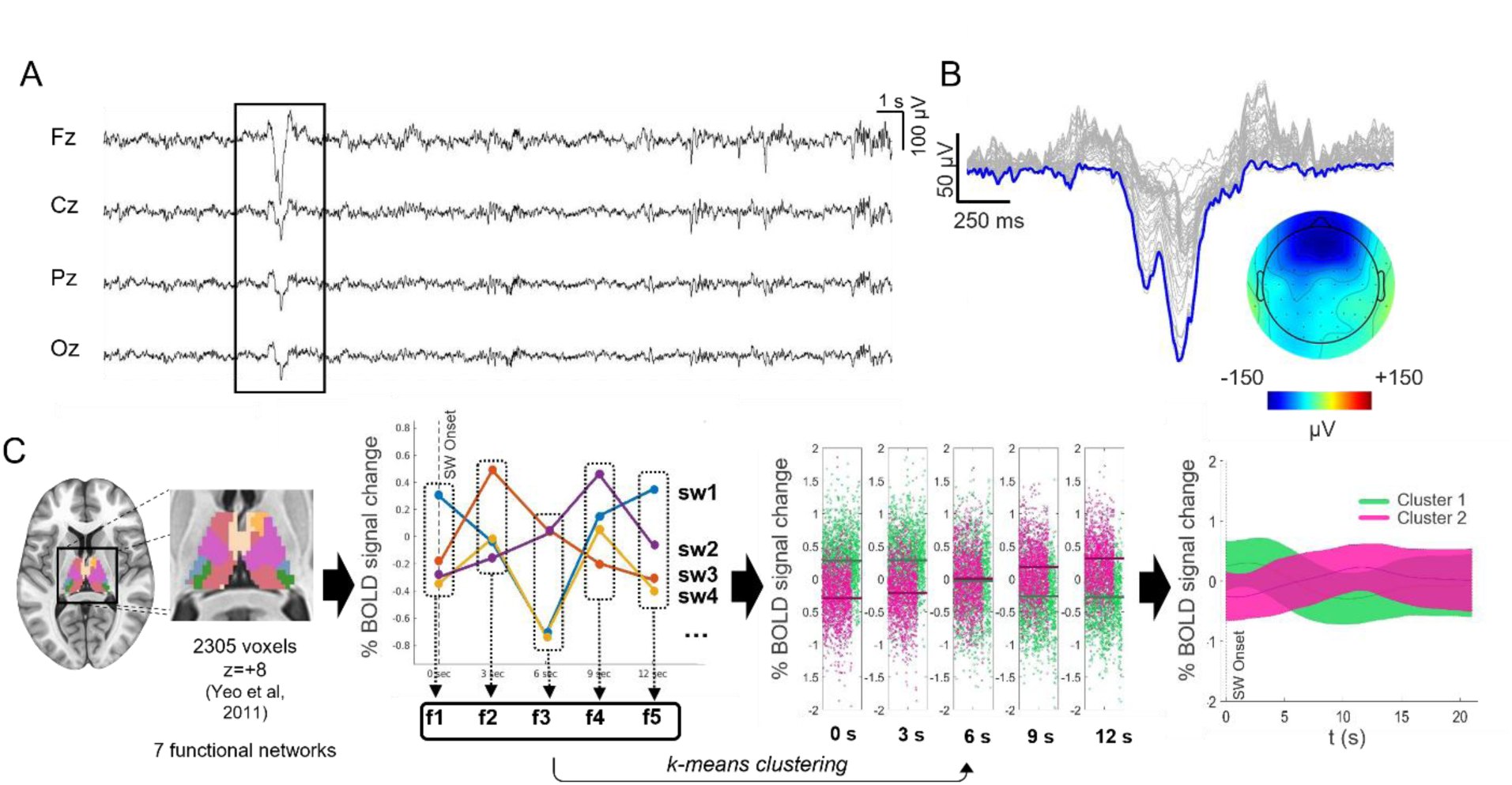
Slow-wave clustering procedure. A. EEG trace of a representative N2 epoch with an example of a detected slow wave. B. The corresponding topographical distribution (slow-wave involvement) at the slow wave negative peak (t=0) is also shown (in µV). C. Diagrammatic illustration of the slow-wave clustering procedure. The average BOLD-signal was extracted from each of seven thalamic functional ROIs. The signals were concatenated. For each identified slow wave and thalamic network, five features were extracted, reflecting the BOLD signal sampled at each TR from 0 to 12 seconds post-slow-wave onset. This resulted in a total of 35 features (5 for each of the 7 ROIs). We employed a k-means clustering approach, using correlation as the measure of distance, to differentiate slow waves according to the selected features. Each slow wave was represented as a 35-dimensional point in the clustering space and was assigned to one of the two clusters based on its distance from the centroid. The bottom-right plot illustrates the mean (± SD) BOLD-signal for two distinct clusters of slow calculated for a single subject in one of the seven functional networks.

**Table 1.**
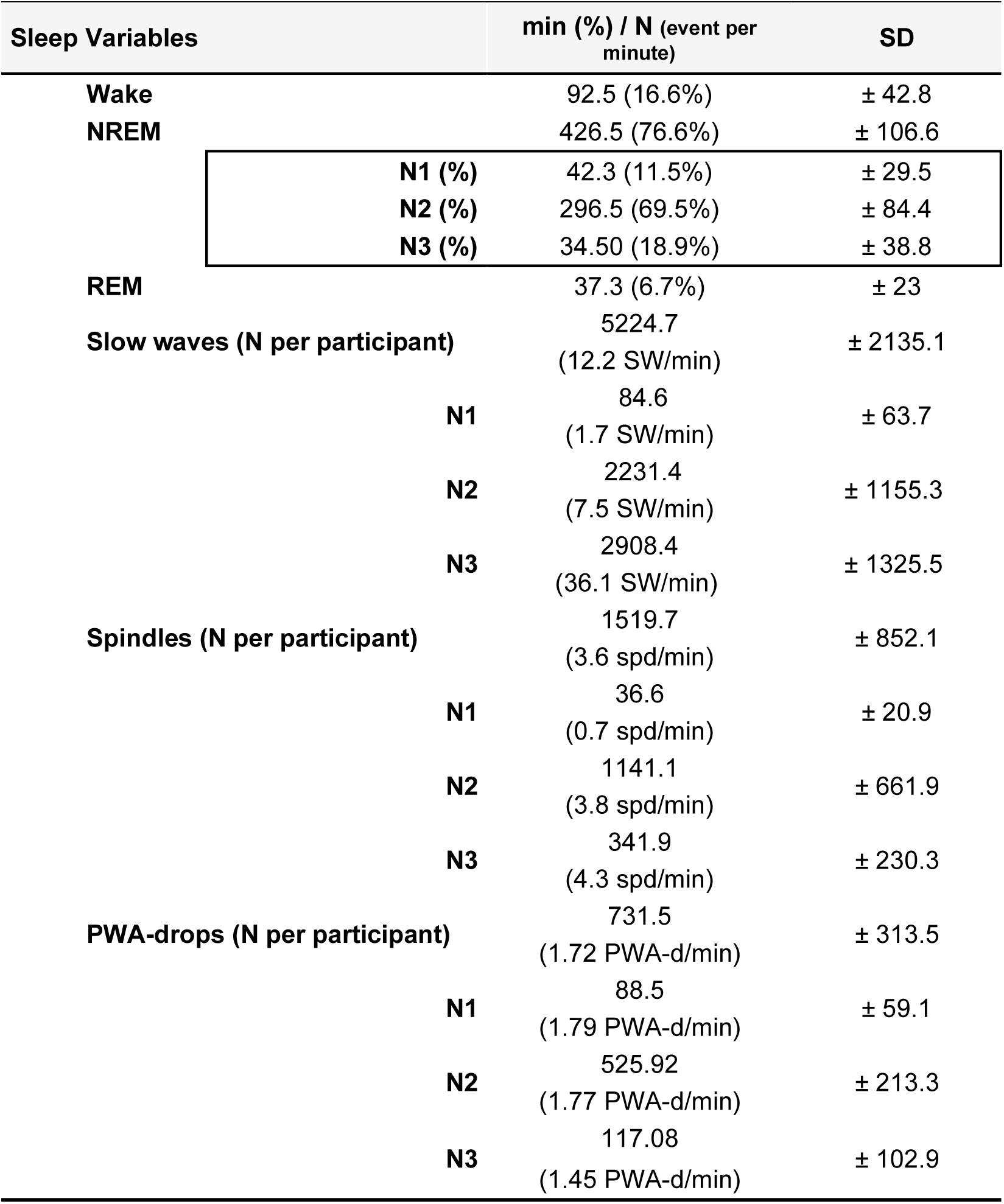
Sleep-related variables. Results are reported as mean ± SD or density/minute. PWA: Pulse Wave Amplitude.

### Two slow-wave types are identified based on thalamic activity

We categorized the EEG slow waves based on their corresponding thalamic fMRI activity using a k-means clustering algorithm (Figure 1). For this purpose, wave-locked BOLD-signals from seven thalamic regions of interest (ROIs) were extracted and concatenated (0–12 s relative to slow wave onset; see Methods for details). These ROIs were defined based on thalamic voxels’ preferential functional connectivity with classical cortical networks^40^. The analysis identified two distinct slow-wave clusters that were highly consistent across participants. Slow waves in Cluster 1 (C1) exhibited an early positive BOLD response across all seven networks, peaking approximately 3 s after wave onset, followed by a negative BOLD change around 10 s post-onset (Figure 2 A). In contrast, slow waves in Cluster 2 (C2) displayed an initial negative BOLD deflection between 3 and 4 s after wave onset, which was followed by a positive BOLD change around 12 s post-onset (Figure 2 A).

**Figure 2.**
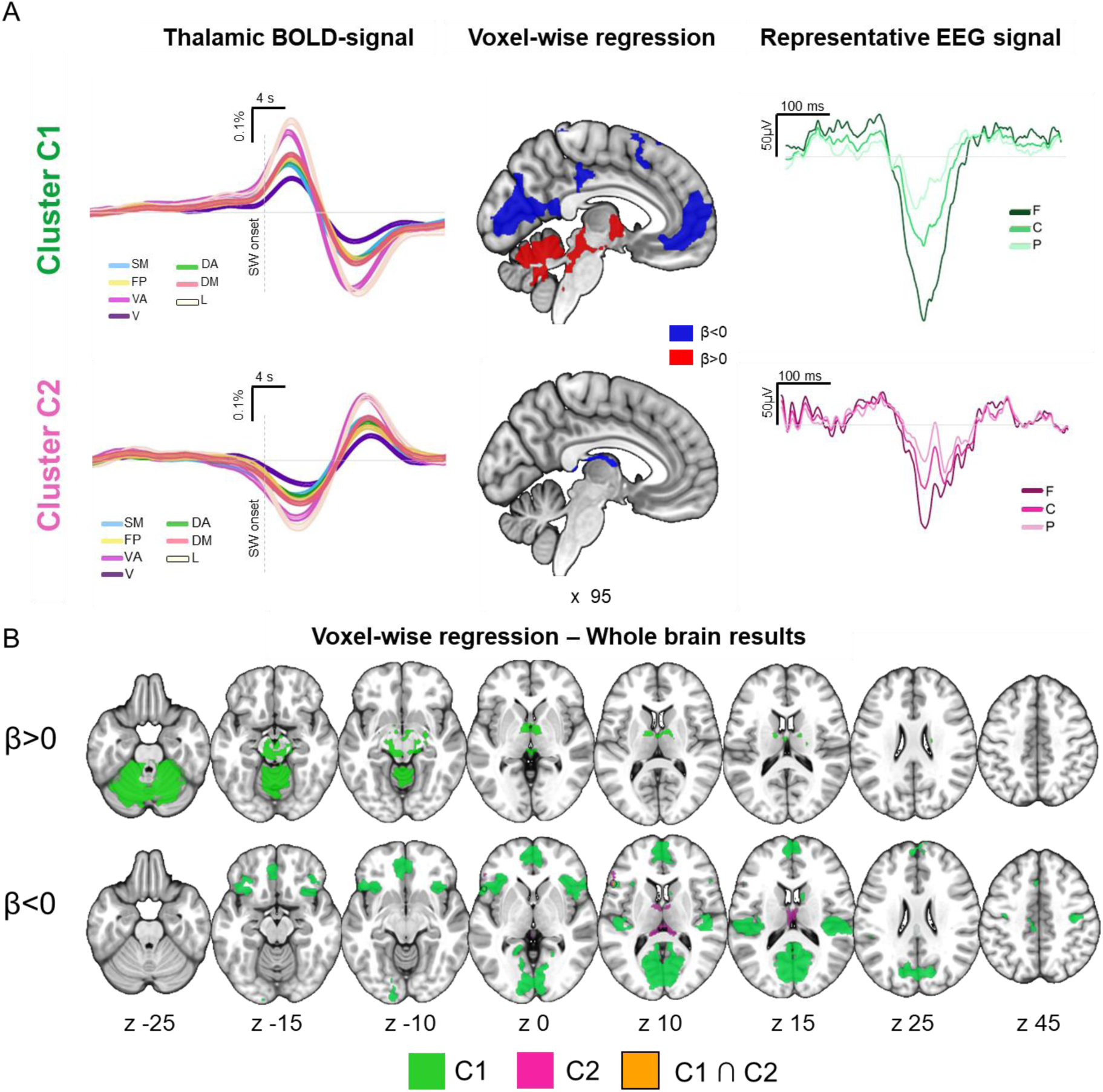
BOLD and EEG signal changes associated with the two slow-wave clusters. A. The left panel displays the mean changes in the BOLD signal (± standard error) time-locked to C1 and C2 slow waves across seven distinct thalamic networks. The BOLD signals were up-sampled to match the EEG sampling rate and aligned with the onset of the slow waves. The central panel shows significant positive (red; >0) and negative (blue; <0) BOLD-signal changes in a medial sagittal view, obtained through a voxel-wise regression analysis in which the occurrence of C1 and C2 slow-waves were modeled as two separate regressors.In the right panel, EEG traces for two representative slow waves from each cluster are shown (±0.5 second window around the negative peak of the slow wave). B. Brain regions (q < 0.001) associated with significant BOLD-signal increases (top; >0) and decreases (bottom; <0) for C1 (green) and C2 (magenta) slow waves. A small overlap area (orange) was found in the right inferior frontal gyrus. Brain images were generated using MRIcroGL (https://www.nitrc.org/projects/mricrogl/). SM: somatomotor; FP: fronto-parietal; VA: ventral attention; v: Visual; DA: dorsal attention; DM: default mode; L: limbic; F: average signal from frontal electrodes (F3, Fz and F4); C: average signal from central electrodes (C3, Cz and C4); P: average signal from parietal electrodes (P3, Pz and P4).

Voxel-wise whole-brain regression conducted including the two clusters revealed distinct patterns of wave-locked hemodynamic changes (Figure 2 B; Supplementary Table S4). C1 slow waves were associated with widespread positive BOLD changes in subcortical regions, including the pons, midbrain, cerebellum, and thalamus, coupled with negative BOLD changes in cortical areas such as the medial frontal cortex, parieto-occipital regions, and somatomotor areas. In contrast, C2 slow waves showed more localized negative BOLD changes, primarily in the medial thalamus and inferior frontal cortex. Notably, the C1 hemodynamic pattern closely resembled stage-related BOLD-signal changes observed during light NREM sleep (N1, N2; Supplementary Figure S5, Table S5), suggesting that C1 slow waves may play a significant role in driving these stage-specific hemodynamic variations.

### Morphology and distribution of slow waves

C1 slow waves exhibited significantly broader scalp involvement (p<0.05, paired t-test with cluster-mass correction), higher amplitude (Wilcoxon paired signed rank test; V = 78, q = 0.002, FDR correction), longer duration (V = 78, q = 0.002), and greater synchronization efficiency (paired t-test; t(11) = 7.504, q < 0.001, FDR correction) compared to C2 slow waves (Figure 3). Additionally, C1 waves were more likely to occur in isolation (t(11) = 5.774, q < 0.001) and had a higher density (number per minute) during light (N2) NREM sleep relative to C2 waves (t(11) = 10.174, q < 0.001). No significant differences in density were observed between the clusters during N1 and N3 sleep. A significantly higher proportion of C1 waves were classified as K-complexes compared to C2 waves (t(11) = 3.41, q = 0.021). C1 slow waves were also found to be more frequently associated with sleep spindles (t(11) = 6.789, q < 0.001), during both light N1-N2 sleep (t(11) = 3.585, q = 0.007) and deep N3 sleep (t(11) = 4.552, q = 0.002).

**Figure 3.**
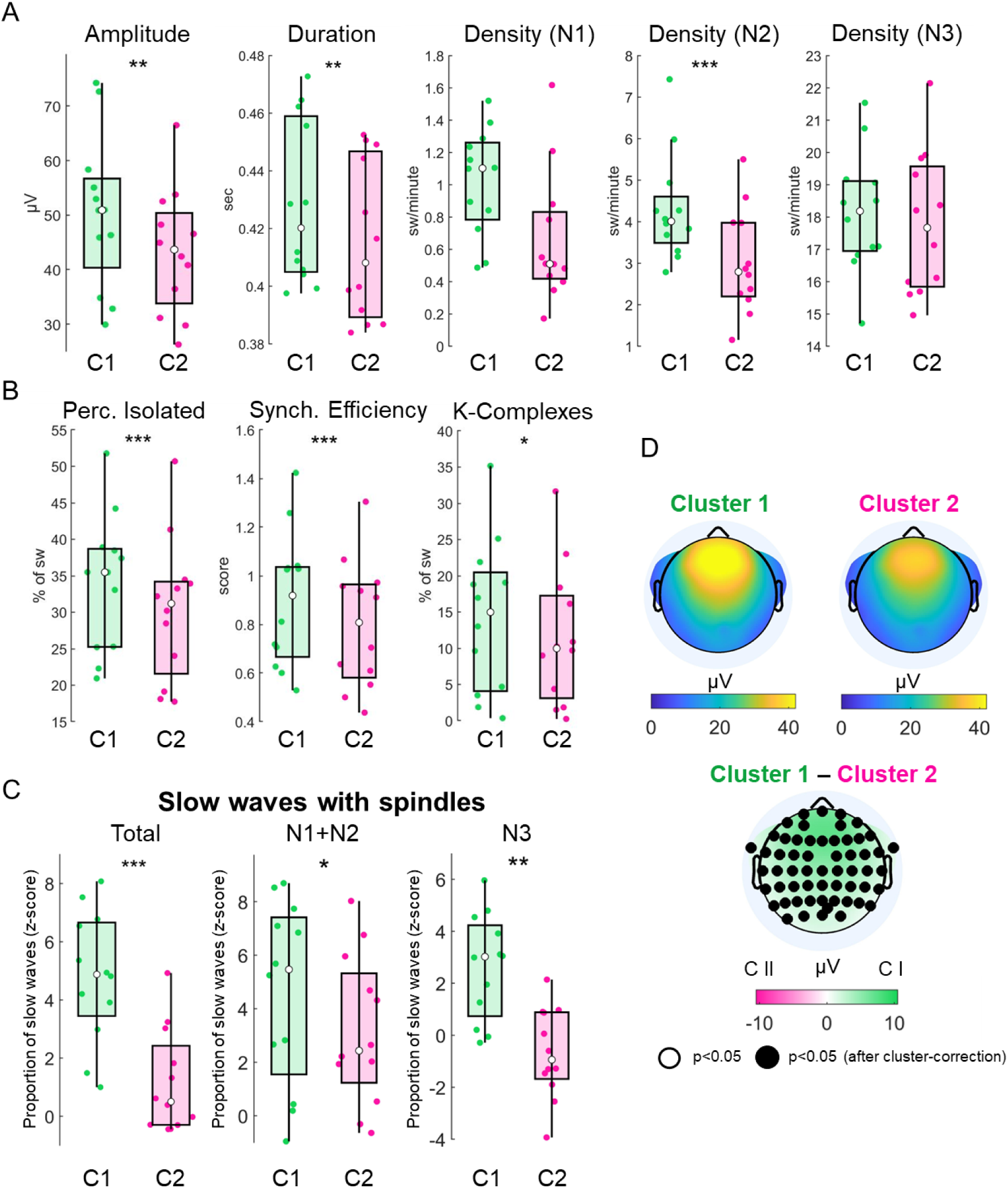
Features and associations of C1 and C2 slow waves. A. Boxplots showing comparisons concerning slow-wave amplitude (µV), duration (s), and density (w/min). B. Boxplots showing comparisons concerning the proportion of slow waves occurring in isolation, slow-wave synchronization efficiency, and the percentage of slow waves identified as K-complexes by a dedicated detection algorithm. In each boxplot, the central dot indicates the median, while the lower and upper edges represent the 25th and 75th percentiles, respectively. C. Z-scored probability of association between slow waves and sleep spindles across all sleep stages (left), in light NREM sleep (middle), and in deep NREM sleep (right). *: q<0.05; **: q<0.01; ***: q<0.001, FDR corrected. D. Absolute scalp involvement of C1 and C2 waves and the statistical comparison between the two. Black dots: p<0.05, cluster-corrected.

### Association between slow waves and sleep fragility periods

During NREM sleep, sigma/spindle activity in centroparietal (somatomotor) brain areas fluctuates at an infraslow timescale around 0.02 Hz (∼50-s cycle ^37^). This oscillation is believed to originate from fluctuations in the activity of the noradrenergic locus coeruleus, which modulates thalamic depolarization and, in turn, spindle generation ^38^. Importantly, these infraslow sigma oscillations are closely linked to variations in sleep stability, defined in terms of arousal/awakening probability in response to external stimuli ^37^. Indeed, the rising phase of the oscillation (140–330°), marked by increased sigma activity, is associated with enhanced sleep stability. In contrast, the descending phase (330–140°), characterized by reduced sigma activity, corresponds to greater sleep fragility and elevated autonomic activity, as reflected by an increase in heart rate^37^. Based on these findings, we sought to investigate potential differences between the two identified slow wave clusters in relation to their coupling phase with the infraslow oscillation. To this aim, we calculated the individualized infraslow peak frequency during N2 sleep, where the sigma power oscillation is most pronounced. Then, we determined the infraslow phase at the onset of each slow wave from the two clusters (including all NREM epochs).

We found that C1 slow waves predominantly occurred during the descending phase of the infraslow oscillation (Figure 4). In contrast, C2 slow waves showed a less defined phase distribution, with higher inter-subject variability (Supplementary Figure S6). Their mean phase of occurrence peaked at ∼140°. In the absence of a package supporting repeated-measures analysis for circular data, we quantified the likelihood of each cluster occurring within the two infraslow phases by computing the probability of occurrence in each phase bin. A repeated-measures (rm)ANOVA revealed a significant interaction between infraslow oscillation phase ^37^ and cluster type (F(1,11): 9534.2, p < 0.001). Post-hoc analysis confirmed that C1 slow waves were significantly more likely to occur during the *descending* phase of the oscillation, corresponding to periods of high sleep fragility (t(11): −6.547, p < 0.001). Conversely, C2 slow waves were more likely to occur during the *rising* phase, associated with periods of low sleep fragility (t(11): 3.332, p = 0.006).

**Figure 4.**
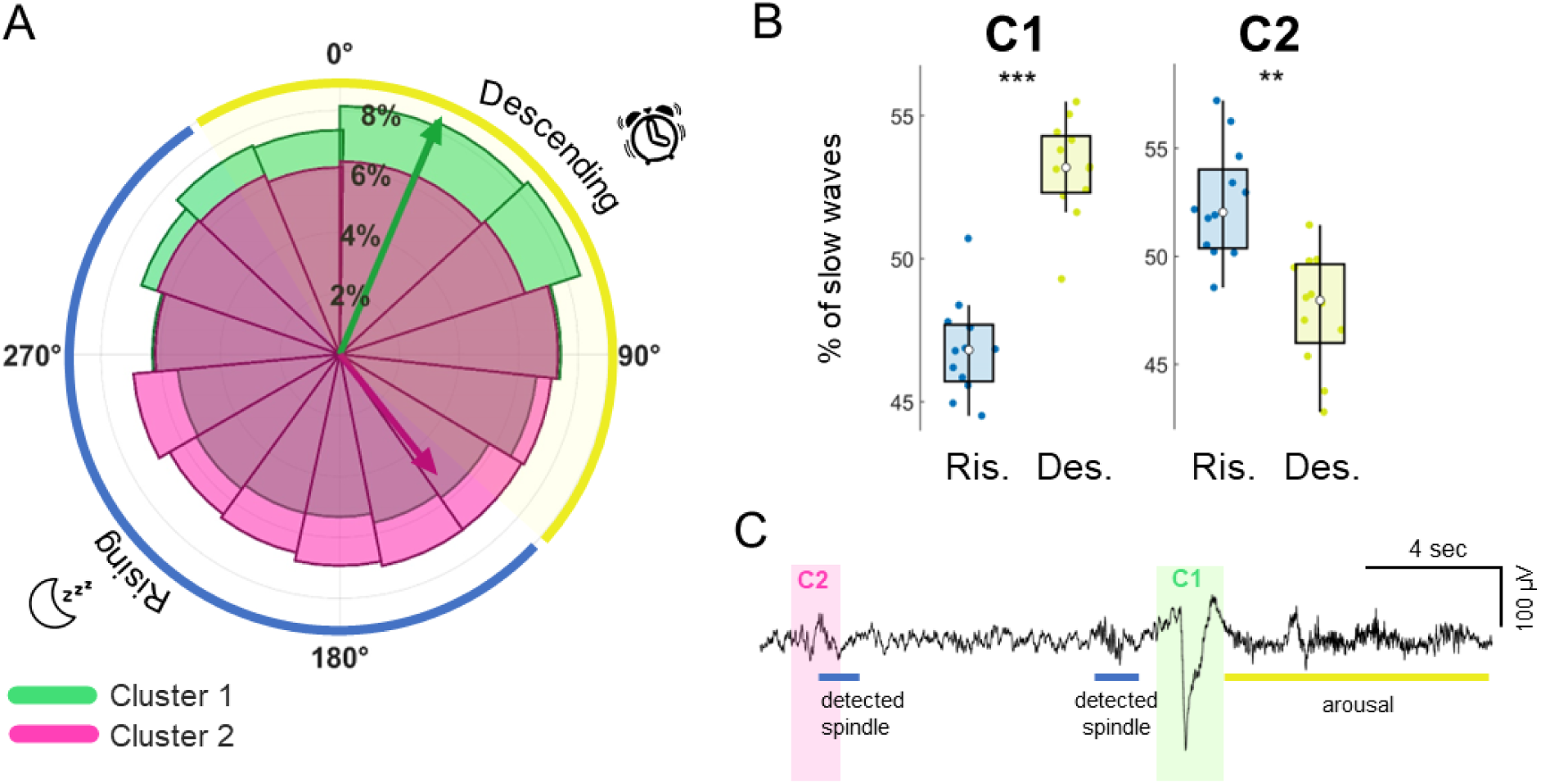
Relationship between slow waves and sigma infraslow oscillation. A. Polar plot showing the mean phase distribution of slow waves with respect to the sigma infraslow oscillation (0° represents the infraslow peak). The full range of the infraslow phase was divided into 15 bins (24° each) for visualization purposes. The yellow-shaded region indicates the descending phase of the infraslow oscillation which corresponds to periods of high sleep fragility. The blue-shaded region represents the rising phase, associated with high sleep stability. B. The boxplots display the percentage of slow waves during the two different infraslow phases, the rising and descending phase, for each cluster. A post-hoc t-test was performed following the application of a repeated-measures ANOVA to assess the interaction cluster by phase. In each boxplot, the central dot indicates the median, while the lower and upper edges represent the 25th and 75th percentiles, respectively. **: p<0.01; ***: p<0.001. C. An example EEG trace marking C1 and C2 slow waves, together with detected spindles and arousal.

### Association between slow waves and peripheral vasoconstriction events

Previous research linked sympathetic activation to fluctuations in the fMRI global signal during sleep, implicating autonomic changes in the modulation of brain-wide hemodynamic responses ^41,42^. These changes could partly arise from vascular mechanisms, such as the redistribution of blood flow across cortical and subcortical structures. According to this perspective, fMRI changes during slow waves - driven by subcortico-cortical, arousal-related synchronizations like K-complexes - may primarily reflect autonomic vascular responses rather than directly resulting from neural activity changes.

Abrupt drops in photoplethysmographic (PPG) pulse wave amplitude (PWA drops) are commonly interpreted as indicators of peripheral vasoconstriction and increased sympathetic activity. This interpretation is supported by evidence from studies showing that PWA drops are tightly associated with cortical EEG arousals, especially during light (N1, N2) NREM sleep ^33,43–45^. Leveraging PWA-drops as indicators of autonomic and arousal system activation, we examined their association with the two identified slow wave clusters. This approach aimed to determine the potential role of autonomic dynamics in modulating cluster-specific slow-wave activity and to explore whether autonomic-mediated mechanisms contribute to BOLD-signal changes linked to slow waves.

First, we investigated the strength of association between slow waves and PWA-drops (Figure 5A). On average, participants exhibited 1,573 ± 1,068 slow waves associated with PWA-drops and 3,652 ± 1,366 waves showing no association with PWA-drops. When analyzing all NREM stages collectively, no significant differences in the likelihood of PWA-drop association were observed across slow-wave clusters. However, a stage-specific analysis revealed distinct patterns. In lighter sleep stages (N1, N2), C1 waves were significantly more likely to coincide with PWA drops compared to C2 waves (t(11) = 3.261, q = 0.02). Conversely, in deeper sleep (N3), the association shifted, with C2 waves being more frequently linked to PWA drops than C1 waves (t(11) = −4.252, q = 0.007; Figure 5B).

**Figure 5.**
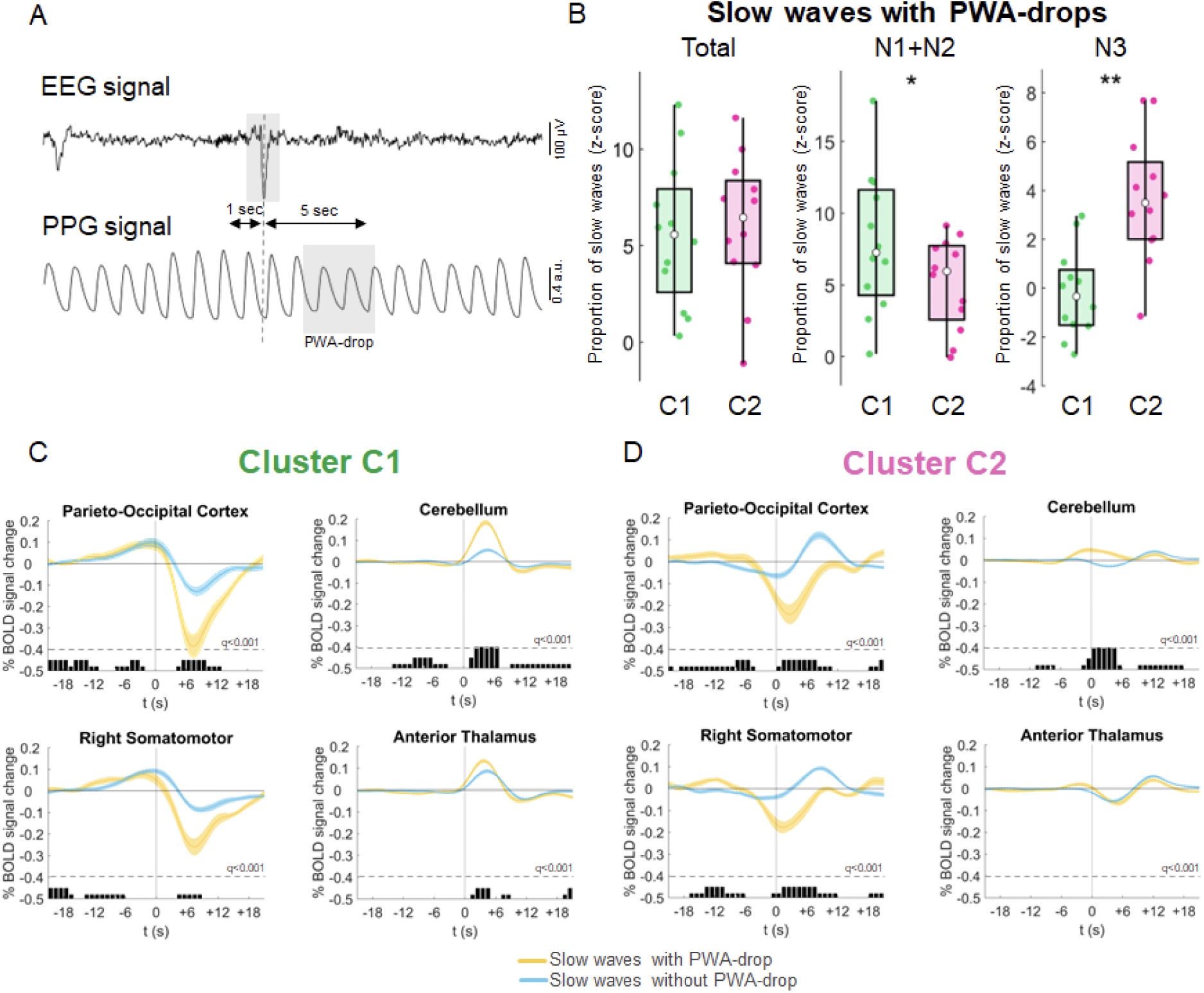
Association between slow waves and PWA drops. A. An example of EEG and photoplethysmographic (PPG) signals, illustrating the temporal relationship between a detected slow wave’s negative peak in the EEG signal (Fz channel, top) and a subsequent drop in the PPG signal (PWA-drop; bottom) within the 5-second temporal window. B. The boxplots display the z-scored probability of association between slow waves and PWA-drops across all sleep stages (left), in light NREM sleep (middle), and in deep NREM sleep (right). In each boxplot, the central dot indicates the median, while the lower and upper edges represent the 25th and 75th percentiles, respectively. *: q<0.05; **: q<0.01; ***: q<0.001, FDR corrected. C. and D. Mean BOLD signal changes time-locked to the onset of Cluster 1 and Cluster 2 slow waves with and without co-occurring PWA-drop are shown alongside their standard errors. These plots compare slow waves associated with a concurrent PWA drop (yellow) to those without a drop (light blue). Significant differences between the two conditions, calculated over 1-second bins and corrected for multiple comparisons using the FDR method, are indicated by gray bars. The BOLD profiles were generated by averaging the signal across individual voxels within significant clusters identified through the main slow wave regression analysis. For representative purposes, only four out of the twelve significant regions of interest (ROIs) derived from the main slow wave regression analysis are illustrated. See Supplementary Figure S4 for the other ROIs. **t₀**: marks the negative peak of the slow wave.

Next, we investigated whether slow waves with and without accompanying PWA drops exhibited distinct response patterns, focusing on key cortical and subcortical regions showing significant hemodynamic changes associated with slow waves (Figure 5C-D also see Supplementary Figure S7). This analysis was performed separately for C1 and C2 waves. Thalamic BOLD responses time-locked to C2 slow waves did not differ significantly between waves with or without associated PWA-drops, while small significant differences were detected for C1 slow waves. However, notable differences emerged in the cerebellum and cortical regions. In the cerebellum, C1 slow waves associated with PWA drops showed a stronger positive BOLD response compared to those without. For C2 waves, an early positive response was present in waves with PWA-drops, whereas a negative-to-positive fluctuation characterized waves without PWA-drops. At the cortical level, PWA-drops during both C1 and C2 waves were linked to pronounced large-amplitude negative BOLD-signal modulations. This modulation was attenuated for C1 waves without PWA-drops. In contrast, for C2 waves, the negative BOLD-signal response nearly disappeared in the absence of PWA-drops, giving way to a delayed positive deflection. Overall, these findings highlight the crucial role of vascular dynamics, as reflected by PWA-drops, in shaping slow-wave-associated hemodynamic responses.

## Discussion

The sleep slow wave, the main hallmark of NREM sleep, plays essential roles in several fundamental processes, including sleep-dependent memory consolidation through the coordination of spindles and hippocampal ripples ^46^, metabolic waste clearance, and sensory disconnection. In humans, slow waves are typically studied using scalp EEG, while their associated changes in subcortical activity have rarely been explored. Investigations in this area primarily relied on intracranial recordings from clinical populations. (e.g., ^18,47^). In this study, we employed simultaneous whole-night EEG-fMRI recordings to examine thalamic activity changes associated with sleep slow waves. Our findings reveal evidence for at least two distinct slow-wave clusters, each characterized by unique functional correlates across cortical and subcortical regions. The first cluster (C1) is characterized by an early positive thalamic response and includes larger, longer, and highly synchronized slow waves. These waves are associated with widespread hemodynamic changes across cortical and subcortical regions. C1 waves are more prevalent during lighter NREM stages (N2), and during sleep periods previously shown to be more sensitive to external stimuli and prone to arousal. Additionally, these waves are strongly linked to sleep spindles and, during light sleep, to autonomic markers of sympathetic activation and arousal. In contrast, the second cluster (C2) is characterized by an early negative followed by a late positive thalamic response and consists of smaller, less synchronized slow waves. These waves tended to more often occur during sleep phases associated with reduced sensory responsiveness. Unlike C1 waves, C2 waves do not exhibit consistent hemodynamic patterns across cortical and subcortical structures, likely due to more variable origins and spatial distribution. These findings suggest that slow waves are not a monolithic phenomenon but encompass distinct functional subtypes with unique roles in sleep physiology. Interestingly, previous work suggested the existence of a three-way association between EEG slow waves, hemodynamic cortical changes, and CSF movements, and suggested that this interaction may ultimately favor the movement of interstitial fluids and the removal of metabolic wastes from brain tissues ^7,8^. Our results suggest that C1 and C2 slow waves, showing a different association with hemodynamic changes, might have a different role in or association with metabolic clearance mechanisms.

The notion that sleep slow waves in the delta frequency range (<4 Hz) may represent distinct processes and fall into separate subclasses has a long history. Early pioneering studies by Steriade and Amzica using animal models made a clear distinction between slow oscillations (<1 Hz) and delta oscillations (1–4 Hz), emphasizing differences in their generation and functional significance ^10,48^. More recent research in animals demonstrated that slower and faster delta-band waves may differ in their homeostatic regulation and roles in memory consolidation (e.g., ^20,27,49^). Indeed, slower waves, as opposed to faster ones, appear to favor a weakening, rather than a consolidation, of recently acquired memories. Moreover, they have been shown to undergo a less pronounced increase following sleep deprivation and a weaker decline over the course of a night. In humans, evidence supporting a similar differentiation has emerged. Indeed, one investigation showed that slower waves peaking below 1 Hz may exhibit a weaker homeostatic decline across a night of sleep compared to faster delta waves ^19^. However, classifying slow waves purely by frequency is challenging. Significant overlap is inevitable between adjacent frequencies, and the peak frequencies of slow wave sub-types may vary across species, individuals, developmental stages, and pathological conditions.

Several studies have proposed alternative classifications of human slow waves based on their temporal, morphological, and topographical characteristics. For instance, an analysis of slow-wave changes during the wake-sleep transition identified two distinct processes underlying slow-wave generation and synchronization: an early process, responsible for producing large-amplitude, widespread, and highly synchronized slow waves on a background of small-amplitude oscillations, and a later process, characterized by the emergence of smaller, more local, and less synchronized slow waves ^3^. The appearance of highly synchronized slow waves -likely including so-called K-complexes-in a phase dominated by poorly synchronized activity is thought to involve diffusely projecting systems, such as subcortical arousal-related structures, which can coordinate widespread slow-wave generation across distributed neuronal populations. In line with this view, early slow waves, referred to as type I waves, tend to mainly originate in the somatomotor cortex, a region heavily innervated by the noradrenergic locus coeruleus. In contrast, late slow waves, also indicated as type II, likely arise from less efficient cortico-cortical or cortico-subcortico-cortical processes. Further research demonstrated that type I and type II waves persist into stable NREM sleep, where they exhibit distinct patterns of homeostatic regulation. Indeed, type II waves, but not type I waves, display a clear decline in amplitude and number throughout the night ^3,21^. Consistent with this, recent work showed that large and widespread slow waves with no homeostatic overnight decline similar to type I waves increase in number following sensory stimulation, while this is not observed for smaller, more local slow waves compatible with type II waves ^21,50^. Developmental studies have also highlighted differences in the trajectories of these two wave types and their synchronization mechanisms. For instance, research in school-aged children and adolescents revealed distinct maturation patterns for type I and type II waves ^51,52^, potentially driven by changes in thalamic contributions to cortical synchronization ^53^.

Our findings are largely consistent with prior observations using scalp EEG, supporting the existence of two distinct slow-wave subtypes with differential thalamic involvement. In this study, we deliberately focused on the thalamus rather than other arousal-related structures previously implicated in slow-wave synchronization. This choice was driven by both technical considerations and pre-existing evidence. Technically, brainstem and basal forebrain regions involved in arousal and sleep regulation commonly exhibit lower signal-to-noise ratios in fMRI data compared to neocortical structures and the thalamus. This is attributed to the small size of their nuclei and the close proximity to blood vessels and CSF flow ^54–56^. These factors make detailed investigations of these regions more complex compared to the thalamus. From an evidence-based perspective, numerous studies in animals and humans have highlighted the critical role of the thalamus in regulating slow-wave dynamics ^14,16,40,47,57,58^. Emerging evidence even suggests that the thalamus may act as a leading driver of cortical slow waves, initiating bottom-up synchronization ^18^. This potential role of the thalamus in driving type I slow waves aligns with its broader functional organization. Beyond its classical role in sensory and motor signal relay, the thalamus is central to arousal modulation across states of wakefulness and sleep ^59,60^. Its connections with the ascending reticular activating system and brainstem nuclei such as the locus coeruleus ^61^ position it as a critical hub capable of influencing vigilance and slow-wave generation ^15,18,62–64^. Although the temporal resolution of fMRI does not allow us to definitively establish whether thalamic activity precedes or follows cortical slow waves, our findings revealed a strong association between large, synchronous (C1) slow waves and thalamic dynamics. Consistent with this interpretation, C1 slow waves exhibited a stronger - and, we hypothesize, earlier - association with thalamus-generated sleep spindles compared to C2 waves. Interestingly, though, rather than entirely excluding a thalamic involvement in C2 waves, our results point to a distinct involvement pattern. One intriguing possibility is that the initial negative thalamic signal deflection during C2 slow waves may reflect the cortico-thalamic transfer of slow-wave activity, followed by increased thalamic activation ^47^. This interpretation would be consistent with previous evidence indicating that type II slow waves are followed, in the few seconds after their occurrence, by a relative increase in spindle activity, while a relative suppression of spindle generation is observed seconds after type I waves ^22,65,66^.

Beyond the thalamic involvement, our findings point to a broader engagement of arousal-related mechanisms in C1 slow waves and highlight the significant contribution of autonomic dynamics to slow-wave-associated BOLD signal changes. Indeed, we found that C1 slow waves occurred preferentially during specific phases of a 0.02 Hz infraslow oscillation in sigma/spindle activity (12–15 Hz), a rhythm previously linked to variations in arousability ^37^ and vasomotion-induced glymphatic activity ^67^. Animal studies have demonstrated that these dynamics correspond to fluctuations in noradrenergic locus coeruleus activity during NREM sleep ^38,68^. The increased occurrence of C1 slow waves during infraslow phases associated with heightened noradrenergic activation supports their connection to arousal dynamics and sleep fragility. Additionally, our results indicate a significant association between C1 slow waves and PWA drops, which signify peripheral vasoconstriction during light NREM sleep. Notably, PWA drops are typically accompanied by transient heart rate increases and can be elicited by noradrenaline infusion ^45^. At the cortical level, PWA drops have been linked to EEG activations and microarousals ^44,69^. For these reasons, they are regarded as a signal of autonomic and arousal-related activation ^36^. Crucially, while we found a stronger association between C2 waves and PWA-drops during N3 sleep, PWA-drops are significantly less numerous during this stage relative to N1/N2 sleep and are less often associated with arousal events. The reasons for these differential couplings between slow waves and PWA-drops across NREM substages is unclear, but could reflect broader variations in the coupling between central and peripheral arousal dynamics.

Notably, PWA drops are associated with a diffuse reduction in cortical BOLD-signal, likely reflecting autonomic-mediated vascular changes in the brain ^41,70^. Given the relationship between PWA-drops and slow waves, such vascular dynamics could contribute to BOLD-signal changes observed during slow waves. Indeed, we found that C1 and C2 slow waves exhibited partially distinct BOLD-signal patterns depending on whether they coincided with PWA-drops. For C1 slow waves, the BOLD dynamics partially overlapped with those of PWA-drops, suggesting that a summation effect might take place in certain brain regions when these events co-occur. In contrast, C2 slow waves displayed markedly different BOLD dynamics depending on the presence or absence of PWA-drops, potentially explaining the lack of consistent BOLD correlates for these waves at cortical and subcortical levels. These observations suggest that BOLD-signal changes during slow waves likely result from a combination of neural and autonomic-mediated vascular dynamics, with the relative contribution of each varying across slow-wave subtypes due to their differing associations with arousal dynamics.

## Limitations

Several limitations of the present study should be acknowledged. First, the scanner environment is relatively uncomfortable and noisy, and this may have affected sleep quality and microstructure. For instance, the presence of the high density of C1 waves could reflect a repeated activation of the arousal system due to scanner noise. Moreover, sleep was frequently interrupted during the night, thus limiting the possibility to explore homeostatic changes in slow-wave characteristics. Second, as discussed above, BOLD signal changes are an indirect measure of neuronal activity, and non-neural changes are also known to potentially cause local or global hemodynamic variations (e.g., due to sympathetic vasoconstriction or CO_2_-mediated vasodilation). As such, the contribution of neural and non-neural changes to observed slow-wave-locked variations remains unclear. Third, slow waves may occur in trains or even near-continuous oscillations during N3 sleep. The occurrence of slow waves in close temporal vicinity may complicate their analysis and classification, especially given the slow temporal evolution of BOLD-signal changes ^71^.

## Conclusions

The present study identified two distinct clusters of NREM slow waves based on different BOLD-signal changes in the thalamus: large amplitude, widespread, and efficiently synchronized C1 slow waves, associated with an early thalamic and subcortical activation, and smaller and shallower C2 slow waves associated with an early negative and a late positive thalamic response. These findings suggest that C1 and C2 slow waves likely arise from different generation and synchronization mechanisms, are subject to distinct regulatory processes, and may fulfill specialized roles in sleep-related brain functions. Furthermore, their differential profiles raise the possibility that they could be selectively impacted in various neurological and psychiatric disorders. Finally, while this study focused exclusively on NREM slow waves, our approach and findings may pave the way for a more accurate characterization and understanding of NREM-like slow waves observed in other states, such as REM sleep or wakefulness ^72,73^.

## Methods

### Participants

We analyzed 24 overnight EEG-fMRI sleep datasets collected over two sessions from 12 healthy adults (8 females, mean age 24 ± 3.46, age range 18-34 years). All data were collected under human subject research protocols at the National Institutes of Health In Vivo NMR Research Center (USA). The Local Institutional Review Board approved the study. The following exclusion criteria were applied during participant screening: any history of neurological illness (e.g., epilepsy), seizures, or prior central nervous system surgery; any current psychiatric diagnosis or lifetime history of major psychiatric illness (e.g., psychosis, bipolar disorder, major depression); uncontrolled medical conditions (e.g., hypertension), pregnancy or nursing, MRI contraindications, or significant hearing impairment; sleep-related disorders including narcolepsy (Type 1), obstructive sleep apnea, chronic insomnia, sleep bruxism, and recurrent sleepwalking in either childhood or adulthood; night-shift or rotating work schedules; excessive stimulant use (caffeine intake >600 mg/day), heavy nicotine or alcohol consumption. Additional exclusion criteria included a score greater than 42 on the Glasgow Content of Thoughts Inventory, a score higher than 10 on the MRI Fear Survey Schedule, a history of chronic back pain, inability to sleep in a supine position and inability to attend a two-night inpatient visit on weekdays throughout the year. An in-person screening was conducted to verify eligibility, and written informed consent was obtained from all participants.

### Experimental Protocol

A detailed description of the study protocol can be found in <u>Moehlman et al., 2019</u>. Participants were scanned twice during two overnight sleep periods separated by a 1-day interval. No sleep deprivation or sedation protocols were used. In order to maximize the probability of sleeping during the MRI scan, participants followed a 14-day sleep hygiene protocol, which included refraining from taking afternoon naps. Adherence to the protocol was verified through actigraphic monitoring. To increase the subject’s comfort inside the scanner, foam pads, and memory foam mattresses were used, while an active acoustic noise reduction system provided hearing protection. Patients were asked to remain still and supine during the overnight scanning procedure. Sleep scans began at 23:00 and ended at approximately 07:00.

Whole-brain fMRI data were recorded with a 3 T Siemens MRI 70 cm bore scanner (Skyra, Siemens, Munich, Germany) and a Siemens 20-channel head coil (TR= 3,000 ms; in-plane matrix = 2.5×2.5 mm^2^; 50 axial slices; slice thickness = 2 mm; slice gap = 0.5 mm; FA= 90°; TE= 36 ms). Data were collected with multi-slice echo-planar imaging in a slice-interleaved fashion. A high-resolution T_1_-weighted anatomical image was also obtained in all subjects (176 sagittal slices; in-plane matrix = 256 × 256; voxel size = 1 × 1 × 1 mm^3^). Polysomnographic data were concurrently collected during the fMRI acquisition using an MR-compatible 64-channel amplifier (BrainAmp, Brain Products GmbH, Germany) including 61 EEG scalp electrodes, 2 EOG electrodes, and 1 ECG electrode. The EEG channels were referenced to the frontal-central midline (FCz) electrode and sampled at 5 kHz. The system received and recorded MRI-scanner triggers for subsequent artifact removal. PPG and chest belt signals were also recorded using a Biopac System (Biopac, Goleta, CA, USA, using TSD200-MRI and TSD221-MRI transducers, an MP 150 digitizer sampling at 1000 Hz using AcqKnowledge software). The PPG signal was recorded from the left index finger.

### Preprocessing of EEG signal

EEG recordings were preprocessed using *BrainVision Analyzer* (Brain Vision, Morrisville, USA) and the *EEGLAB* toolbox ^74^; version 2021.0) for *MATLAB* (The MathWorks, Inc, version 2021a). In particular, the gradient and cardio-ballistic artifact corrections were implemented using standard routines in *BrainVision Analyzer*. For the removal of gradient artifacts, an average artifact subtraction technique was used ^75^. Then, the signal was downsampled to a frequency of 250 Hz. Cardio-ballistic artifacts were removed through a two-step procedure: a template subtraction method was applied, followed by the elimination of artifact-related components from the EEG data using an inverse Independent Component Analysis (ICA) approach ^76^. In order to perform sleep staging, we created a copy of the partially preprocessed EEG signal that was re-referenced to the average of mastoid channels and band-pass filtered using a first-order FIR filter (Hamming type) between 0.3 and 18 Hz. Sleep scoring was then conducted over 30-s epochs according to standard criteria ^1^.

Further preprocessing steps were implemented in *MATLAB* and *EEGLAB* following a standardized procedure described in previous work ^40,53^. First, we created a copy of the partially preprocessed EEG data and used it to identify potential bad channels. This data was re-referenced to the average of all electrodes and high-pass filtered at 1 Hz. Then, bad channels were identified using the *findNoisyChannels* function available within the *PREP-pipeline* plugin ^77^. This approach uses multiple criteria, such as deviation, correlation, predictability, and noisiness to identify bad channels. Then, all runs obtained during the same nocturnal session were merged, and an ICA-based procedure was used to reduce residual EEG artifacts. To this purpose, a copy of the partially preprocessed EEG signal was high-pass filtered at 0.3 Hz and a low pass filtered at 45 Hz, and good channels and NREM epochs free of manually identified artifacts were entered into the ICA routine ^74^. Of note, manually marked artifacts included large residual fMRI- and movement-related artifacts. The *IClabel* plugin ^78^ was used to identify artifactual ICs. We discarded all components for which the probability associated with the ‘brain’ label was below an empirically determined threshold of 0.6. This threshold, though arbitrary, was determined to be optimal for balancing signal and noise based on visual inspection. Finally, the weights obtained from the ICA procedure were applied to the unfiltered, uncut (continuous) data to remove artifactual ICs. Bad channels were interpolated using spherical splines from the activity of the nearest electrodes.

### Detection of slow waves and spindles

Slow waves were detected using a validated algorithm based on the identification of signal zero-crossings in an artificial EEG channel representing the negative signal envelope computed across all available electrodes ^3^. This method allows for the identification of both local and widespread slow waves and defines a unique time reference (across electrodes) for each negative wave ^79^. In particular, the negative-going envelope of the 0.3-45 Hz filtered signal was calculated for each time-point by selecting the four most negative samples (across electrodes), discarding the single most negative sample, and taking the average of remaining values. This approach was used to minimize the potential impact of any residual large-amplitude artifactual activity in isolated electrodes. The resulting signal was baseline corrected (zero-mean centered) before applying a negative-half-wave detection based on the identification of consecutive signal zero-crossings ^3,80^. Only negative half-waves detected during NREM sleep epochs (N1/N2/N3) and with a duration comprised between 0.25 s and 1.0 s (full-wave period 0.5–2.0 s, corresponding to a 0.5–2.0 Hz frequency range) were selected for subsequent analyses. No amplitude thresholds were applied^3,22,40,81^.

Since sleep spindles often occur in association with slow waves, we assessed their possible impact on hemodynamic signal changes during slow waves. As previously described, spindles were automatically detected in the signal of channel Cz using a validated algorithm ^82,83^. Specifically, a wavelet-based filter (10–16 Hz) was applied to the EEG time-series using a *b-spline* wavelet ^84^, and the time course of power of the resulting signal was measured by squaring the values and smoothing the time-series using a sliding window of 100 ms. Then, potential spindles were defined as points in which power values passed a high-threshold corresponding to the median plus 4 times the median absolute deviation of signal power. The actual start and end points of identified events were then measured using the crossing times at a second, low-threshold corresponding to the median power plus 2 median absolute deviations. The thresholds were re-computed for each 30-s epoch. Only events with duration between 0.3 and 3 s and detected during N2/N3 sleep or transitional N1 epochs (i.e., N1 epochs adjacent to N2/N3 epochs) were retained for further analyses ^40^. Finally, a power-ratio threshold was applied in order to ensure some specificity of the transient power increases within the spindle range. Specifically, for each potential spindle, we computed the ratio of the mean power in the spindle range (10-16 Hz) over the mean power in the neighboring ranges (8–10 Hz and 16–18 Hz) and we eventually retained the detected event if the obtained value was greater than 3.

### Detection of pulse-wave amplitude drops

The PWA derived from the PPG signal reflects changes in the pulsatile blood volume at the fingertip during each cardiac cycle. The PWA is defined for each cardiac cycle as the difference between the maximum and minimum values of the corresponding blood volume pulse wave. Thus, drops in PWA are commonly assumed to reflect peripheral vasoconstriction and can serve as an indirect measure of sympathetic nervous system activation ^32,43,44^. A validated algorithm was used to automatically identify PWA-drops in the PPG signal ^85^. The algorithm was applied to the PPG signal after the application of a discrete-time filter cutting frequencies below 5 Hz and above 16 Hz. Prior to PWA-signal extraction, the PPG timeseries was smoothed (Savitzky-Golay filter), and constant and linear trends were removed. Firstly, time-points within the PPG waveform that lacked clear consecutive maximum and minimum points were identified and excluded from further consideration. Time-points deviating from plausible heart-rate values were also excluded (threshold = 250 heartbeats/min). The PWA-signal was then smoothed using a moving average filter to improve the signal-to-noise ratio (5 heart-beat window). Local variance and first derivative criteria were employed to identify potential PWA-drops. A ‘baseline mask’ was computed by including all the ‘stable’ PWA-signal tracts lasting at least 2 consecutive heartbeats. Candidate time-points corresponding to potential PWA-drops were defined as local peaks in the time-course of the PWA-local-variance that simultaneously displayed correspondent negative values in the first-derivative estimates. These candidates were retained if they had an amplitude larger than 10% relative to the baseline value, defined as the mean of the closest previous 5 PWA timepoints belonging to the baseline mask. For each confirmed PWA-drop, we estimated its duration and amplitude.

### (f)MRI Data Preprocessing

Preprocessing of (f)MRI data was performed using AFNI (version 22.1.10) 86, FMRIB Software Library (FSL, version 6.0.5.1) 87, and Advanced Normalization Tools (ANTs, http://stnava.github.io/ANTs/, version 2.3.5) 88.

Anatomical T1w images were skull-stripped with ANTS, aligned to match the EPI (align_epi_anat.py), and non-linearly warped (3dQwarp) using the MNI ICBM152 6th generation atlas 89.

For each fMRI run, slice timing was adjusted to a common temporal origin using 3dTshift. Registration to the first volume was applied for motion correction (3dvolreg). Furthermore, identification of timepoints corrupted by substantial movements was performed using fsl_motion_outliers. Then, a spatial smoothing procedure (3dBlurToFWHM) was applied employing a 7-mm full-width-at-half-maximum (FWHM) Gaussian kernel. Next, for each voxel, we calculated the percent BOLD-signal change relative to the signal mean. The data was non-linearly transformed (3dNwarpApply) for alignment with the ICBM152 atlas coordinate system, and resampled to a 2-mm iso-voxel resolution. To reduce the possible impact of physiological artifacts, activity patterns associated with CSF regions were identified through Principal Component Analysis (PCA). A threshold of 2% was established, such that only those PCs that accounted for at least 2% of the total variance in the CSF signal were retained for subsequent analysis. Finally, regression-based procedures were employed to account for various sources of artifactual activity in the BOLD-signal of each voxel (3dDeconvolve), including head-motion, movement spikes (framewise displacement above 0.3), and CSF-PCs. We also applied temporal autoregression to further account for physiological artifacts (ARMA-1, 3dREMLfit). All runs from the same overnight session were concatenated.

### fMRI-based clustering of EEG slow waves

Slow-wave clusterization was performed based on slow-wave-locked thalamic BOLD-signal profiles extracted from seven ROIs defined according to the preferential connectivity of each voxel with classical functional networks ^40,90^. The decision to use this specific merged configuration of ROIs was based on the assumption that a whole-thalamus mask would fail to capture the distinct functional contributions of individual thalamic regions, which are differentially connected to cortical areas^40^. Before clustering, the BOLD-signal was preprocessed to remove the estimated contribution of the spindle-based regressor, minimizing the potential confounding influence of spindle-activity on the clustering process. Subsequently, for each participant and for every detected slow wave, the mean BOLD-signal profiles across the seven ROIs were calculated and used as input for the k-means clustering procedure. Specifically, we selected a time window of ± 21 seconds around the slow-wave onset, we detrended the signal and used a 12-second time window starting at slow-wave onset for the clusterization procedure. The window-length was chosen according to the typical duration of the BOLD response. The optimal number of clusters was determined using the silhouette criterion ^91^. For each ROI and for each slow wave we extracted 5 features, corresponding to the BOLD-signal value at each fMRI time-point from 0 to 12 seconds after slow wave onset, for a total of 35 features. We used correlation as the measure of distance. Thus, each slow wave was represented as a 35-dimensional point in the clustering space and was assigned to each cluster based on its distance to the corresponding centroid. The clustering was performed at the participant level, followed by mean pairwise correlation similarity calculations to match clusters across participants.

While our chosen clustering configuration proved optimal for the current fMRI dataset, we assessed the robustness of our findings by testing multiple alternative clustering solutions. Specifically, we employed different distance metrics (e.g., cosine instead of correlation) and criteria for determining the optimal number of clusters (e.g., Calinski-Harabasz index instead of the silhouette criterion). We also repeated the clustering procedure using different sets of ROIs, including the limbic thalamus alone, the whole thalamus, the locus coeruleus^85,92^, and a combined set comprising both thalamic and locus coeruleus ROIs. While some of these variations resulted in moderate percentages of slow wave agreement across clustering outcomes, the key properties of the identified slow wave types and their associated thalamic BOLD responses remained highly consistent to the ones shown here. These findings indicate a high degree of robustness and reliability in the clustering of slow waves across methodological and anatomical variations.

### EEG-fMRI regression analysis of slow wave cluster

For each participant, we performed a main voxel-wise regression analysis on BOLD timeseries, using slow waves from the two identified clusters and spindles as regressors of interest. Specifically, spindles were included in our model to disentangle slow-wave-specific BOLD-signal changes from those caused by their common associations with thalamic spindles. We did not include PWA-drops in the model as their possible association with slow waves is less understood. However, we performed additional analyses to evaluate differences across slow waves associated or not with PWA-drops (see below). Slow waves were modeled as square waves with onset corresponding to the first wave zero-crossing, duration equal to the duration of the wave descending phase (from the first zero-crossing to the maximum negative peak), and amplitude corresponding to the absolute value of the maximum negative peak. We included in the regressor all detections in N1, N2, and N3 sleep. Spindles were modeled as squared waves with onset and duration derived from the automated detection algorithm, and fixed amplitude of one. We included in the regressor spindles detected during N2, N3, and N1 epochs adjacent to N2 or N3 epochs.

All the regressors were convolved with a standard gamma hemodynamic response function and down-sampled to match the BOLD time-series sampling rate. Regressors from different runs within the same session were concatenated. For segments containing artifacts, wakefulness periods, or REM sleep, all regressors were set to zero, and BOLD time-series were set to baseline values. We converted beta-values of cortical voxels into z-scores computed with respect to a null distribution of beta-values obtained by shuffling the timing of events of interest (i.e., slow waves, spindles). We ensured that the total number of events remained constant between the original and shuffled regressors and we prevented the repositioning of events within zero-valued data segments.

At the group level, we employed a mixed-effect model to evaluate the statistical significance at each voxel. We treated z-scored beta values as the response variable, while participants were included as a random factor. To account for multiple comparisons, we applied a False-Discovery-Rate (FDR) correction (q < 0.001) ^93^.

### Comparison of slow-wave features across clusters

Slow-wave clusters were compared with respect to several characteristics, including density (number of waves per minute), amplitude, duration of the half-wave, probability of occurrence across NREM substages (N1, N2, N3), probability of occurrence in isolation (no other slow waves peaking within a temporal window of −12 to +5 seconds around slow-wave onset), and synchronization efficiency (synchronization score). As described in previous work, a synchronization score was computed for each wave as the mean of descending and ascending slopes multiplied by the proportion of involved channels showing a local signal below −5 μV ^22^. We also evaluated how many slow waves morphologically defined as K-complexes were found in each cluster. To this aim, K-complexes were detected using a convex optimization algorithm implementing a Teager–Kaiser Energy Operator method (*DETOKS*, ^94^). The algorithm specifically identifies K-complex waves on the envelope of the band-pass filtered signal. Prior to applying the algorithm, the signal was downsampled to 100 Hz and filtered with a high-pass filter at 0.5 Hz and a low-pass filter at 45 Hz. Then, we identified slow waves that corresponded to putative K-complexes by assessing their temporal overlap with events detected by the *DETOKS* algorithm. The percentage of slow waves corresponding to putative K-complexes was calculated and compared between clusters. All comparisons were conducted using paired t-tests for normally distributed data and Wilcoxon signed-rank tests for non-normally distributed data. All p-values were adjusted for multiple comparisons using FDR correction ^93^.

Additionally, we performed topographical comparisons of involvement across the two slow-wave clusters. Slow-wave involvement was defined as the average EEG voltage calculated within a 40-ms time window around the wave negative peak, averaged within subjects. Topographical statistical comparisons across slow-wave clusters were performed using paired t-tests and a cluster-mass correction for multiple comparisons (p < 0.05 cluster forming threshold; N = 10,000 permutations).

### Relationship between slow waves and sigma infraslow oscillation

Previous work demonstrated that sigma power (12-15 Hz) oscillates at an infraslow (0.02 Hz) frequency and that the phase of such an oscillation is related to changes in arousability. Here, we assessed the potential association of slow waves in each identified cluster with specific phases of the infraslow oscillation. We focused this analysis on N2 sleep, as this stage exhibits the most pronounced oscillation in sigma power. EEG signals from the Pz channel were bandpass-filtered in the sigma range using a second-order Butterworth filter. Sigma power was extracted via the Hilbert transform and normalized in decibels (dB). A continuous wavelet transform (CWT) was then applied to the normalized sigma power signal to perform time-frequency decomposition, calculated separately for each sleep stage. Then, the specific infraslow oscillation frequency of each participant was defined as the spectral oscillation peak within the 0.015–0.025 Hz range during N2 sleep. If no clear peak was detected, a default frequency of 0.02 Hz was assigned. Following this, sigma power was re-extracted and normalized for all NREM epochs. Then, the signal was bandpass-filtered around the participant-specific infraslow frequency (±0.005 Hz; second-order Butterworth filter). The Hilbert-transformed phase of the infraslow oscillation was then extracted at the onset of each slow wave. Based on previous findings, we defined two main infraslow phases of interest ^37^: the *descending* phase (330°–140°), indicating higher sleep fragility, and the *rising* phase (140°–330°), reflecting lower sleep fragility. We then calculated the probability of observing C1 or C2 slow waves within each infraslow phase. A repeated measures (rm)ANOVA was used to evaluate the interaction between slow-wave cluster (C1, C2) and infraslow phase (rising, descending). Post-hoc t-tests were performed to explore differences between infraslow phases within each slow wave cluster.

### Associations between clusters’ slow waves and other events

We assessed the potential association of slow waves in each cluster with sleep spindles and PWA-drops. Slow waves were classified as associated with a PWA-drop if the drop occurred between 1 s before and 5 s after the maximum wave negative peak. Instead, slow waves were considered associated with sleep spindles if the spindle occurred within ± 1.2 s with respect to the wave negative peak. For each participant, we computed the probability of slow waves from the two clusters being associated with PWA-drops or spindles. A normalization procedure was applied to ensure comparability of association probability values across slow-wave clusters and NREM substages. Specifically, probabilities were standardized into z-scores relative to a null distribution generated by shuffling event timings (PWA-drops or spindles) within their respective artifact-free sleep-stage epochs (N = 1,000). At the group level, paired t-tests were used to assess differences in z-scored co-occurrence probabilities between the two slow-wave clusters, both for all NREM stages combined and separately for light (N1/N2) and deep (N3) NREM sleep. For both PWA-drops and spindles, p-values were adjusted for multiple comparisons using the FDR correction ^93^.

### Statistical analysis

For all the statistical tests, we used MATLAB (MathWorks). First, we tested for normality of the datasets using the Shapiro-Wilk normality test. For comparisons of two parametric datasets, we used a paired Student’s t test while the equivalent Wilcoxon signed rank test has been used for non-parametric datasets. For comparisons on multiple (> 2) groups of data (as in the case of the interaction between clusters and phase), a repeated measures ANOVA was used (rmANOVA). All the circular data have been analyzed through the CircStat toolbox for MATLAB. Group-level regression analysis on the fMRI BOLD signal was performed using a mixed-effect model (using subjects as random variable). If not stated otherwise, when multiple tests were performed on related hypotheses a False-Discovery-Rate (FDR) adjustment was applied to ensure correction for multiple comparisons.

## Acknowledgments

This work was in part supported by the European Union – Next Generation EU, Innovation Ecosystem “THE - Tuscany Health Ecosystem” (Mission 4, Component 2, Inv. 1.5; code ECS00000017; CUP: D63C22000400001) and by the intramural research program of the National Institute of Neurological Disorders and Stroke, National Institutes of Health.

## Competing Interests

The authors declare no competing interests.

## Data Availability Statement

Source data underlying the results and figures included in the present work are publicly available at this link: https://doi.org/10.18112/openneuro.ds005127.v1.0.4.

## Code Availability Statement

The codes used to preprocess and analyze the data are available upon request from the corresponding authors.

## Notes

https://doi.org/10.18112/openneuro.ds005127.v1.0.4

### Competing Interest Statement

The authors have declared no competing interest.

### Summary of Updates

There was an error in the previous file

