## Supplementary material for "Thalamic involvement defines distinct slow-wave subtypes in NREM sleep"

**Table S1**

| SLOW WAVES | | | | | |
|---|---|---|---|---|---|
| **Signal Change** | **Region** | **Voxels** | **CM x** | **CM y** | **CM z** |
| Positive | Cerebellum | 6107 | 0.8 | -54.9 | -24.2 |
| | Anterior Thalamus | 166 | -0.2 | -3.1 | 0.6 |
| | Left Cerebellum | 106 | -25.6 | -50.7 | -49.6 |
| Negative | Parieto-Occipital Cortex | 4392 | -1.3 | -67.8 | 14.1 |
| | Left Inferior Frontal Gyrus | 1790 | -43.1 | 22.3 | -1 |
| | Medial Superior Frontal Gyrus | 1790 | -1.6 | 54.6 | 3.7 |
| | Right Inferior Frontal Gyrus | 1702 | 41.7 | 23.4 | -3.7 |
| | Right Superior Temporal Gyrus (Heschel Gyrus) | 853 | 47.4 | -25.1 | 14 |
| | Left Superior Temporal Gyrus (Heschel Gyrus) | 830 | 48 | -25.4 | 14 |
| | Medial Superior Frontal Gyrus | 554 | -0.1 | 20.3 | 52.8 |
| | Right Somatomotor | 141 | 40.1 | -18 | 48.5 |
| | Left Somatomotor | 69 | -42.1 | -20.5 | 53.6 |

***Table S1. Brain areas showing significant BOLD-signal changes time-locked to the occurrence of sleep slow waves.*** *The table includes for each area the number of voxels and the coordinates of the center of mass (CM) in the standard MNI space (minimum cluster size = 50 voxels).*

**Table S2**

| SLEEP SPINDLES | | | | | |
|---|---|---|---|---|---|
| **Signal Change** | **Region** | **Voxels** | **CM x** | **CM y** | **CM z** |
| Positive | Insula-Hippocampus-Thalamus | 6972 | -15.2 | -6.9 | 2.9 |
| | Medial Frontal Cortex | 2960 | 0.5 | 12.5 | 43.7 |
| | Inferior Occipital Gyrus | 2526 | 11.9 | -90.3 | -8.6 |
| | Right Somatomotor Cortex | 468 | 52 | -4.3 | 40.7 |
| | Right Inferior Frontal Gyrus | 276 | 51.2 | 15.6 | -6.6 |
| | Left Fusiform Gyrus | 215 | -46.3 | -53.6 | -18.7 |
| | Right Medial Frontal Cortex | 141 | 5.8 | 53 | 18.7 |
| | Right Middle Temporal Gyrus | 124 | 51.7 | -39 | 5.2 |
| | Left Supramarginal Gyrus | 95 | -62.8 | -50.1 | 27.3 |
| | Left Superior Temporal Gyrus | 78 | -56.5 | -28.4 | 0.6 |

***Table S2. Brain areas showing a significant BOLD-signal change time-locked to the occurrence of sleep spindles.*** *The table includes for each area the number of voxels and the coordinates of the center of mass (CM) in the standard MNI space (minimum cluster size = 50 voxels).*

**Table S3**

| PWA-drops | | | | | |
|---|---|---|---|---|---|
| **Signal Change** | **Region** | **Voxels** | **CM x** | **CM y** | **CM z** |
| Positive | Right Caudate | 203 | 14.3 | 5 | 15.2 |
| | Left Caudate | 150 | -15.3 | 5.5 | 15.5 |
| Negative | Roi 1 | 127108 | | | |
| | Left Cerebellum | 567 | -21.5 | -52.4 | -51.8 |
| | Right Cerebllum | 434 | 22.4 | -49.5 | -52 |
| | Cerebellar Vermis | 72 | -1.5 | -71.6 | -34.4 |

***Table S3. Brain areas showing significant BOLD-signal changes time-locked to the occurrence of PWA-drops.*** *The table includes for each area the number of voxels and the coordinates of the center of mass (CM) in the standard MNI space (minimum cluster size = 50 voxels). Roi 1 encompasses nearly all cortical areas, as well as specific subcortical structures, namely the thalamus and pons.*

**Table S4**

| CLUSTER 1 SLOW WAVES | | | | | |
|---|---|---|---|---|---|
| **Signal Change** | **Region** | **Voxels** | **CM x** | **CM y** | **CM z** |
| Positive | Cerebellum, Thalamus, Midbrain, Pons | 8016 | -0.6 | -49.6 | -22.8 |
| | Right Cerebellum | 94 | 12.7 | -67.3 | -45.1 |
| Negative | Occipital Cortex | 4546 | -1.4 | -74.1 | 11.6 |
| | Medial Frontal Cortex | 1736 | -1.5 | 53.1 | 2.4 |
| | Left Inferior Frontal Gyrus | 1088 | -40.5 | 23 | -4 |
| | Right Inferior Frontal Gyrus | 995 | 40 | 23 | -4.3 |
| | Right Superior Temporal Gyrus (Heschel Gyrus) | 783 | 46.6 | 25.8 | 15 |
| | Left Superior Temporal Gyrus (Heschel Gyrus) | 692 | -46.6 | -26.3 | 14.9 |
| | Right Somatomotor | 286 | 30 | -19.7 | 49.6 |
| | Left Somatomotor | 239 | -42.8 | -18.8 | 56.7 |
| | Posterior Cingulate Cortex | 236 | -2.4 | -27.7 | 35.5 |
| | Medial Frontal Cortex | 186 | -2.1 | 14.2 | 60.1 |
| | Somatomotor | 53 | -0.9 | -42.7 | 71.8 |
| | Caudate Nucleus | 51 | 10.2 | 7.2 | 13.4 |

| CLUSTER 2 SLOW WAVES | | | | | |
|---|---|---|---|---|---|
| **Signal Change** | **Region** | **Voxels** | **CM x** | **CM y** | **CM z** |
| Negative | Thalamus | 382 | -0.5 | -24.6 | 9.5 |

| | | | | | |
|---|---|---|---|---|---|
| | Left Inferior Frontal Gyrus | 113 | -53 | 29 | 9.3 |

***Table S4. Brain areas showing significant BOLD-signal changes time-locked to the occurrence of Cluster 1 and Cluster 2 slow waves.*** *The table includes for each area the number of voxels and the coordinates of the center of mass (CM) in the standard MNI space (minimum cluster size = 50 voxels).*

**Table S5**

| N1-N2 SLOW WAVES | | | | | |
|---|---|---|---|---|---|
| **Signal Change** | **Region** | **Voxels** | **CM x** | **CM y** | **CM z** |
| Positive | Cerebellum, Thalamus, Midbrain, Pons | 14680 | -0,4 | -54 | -28,6 |
| Negative | Medial Frontal Cortex | 4052 | -0,8 | 40,5 | 22,4 |
| | Left Insula | 3824 | -45,5 | 3,8 | 7,4 |
| | Bilateral Visual Cortex | 3680 | -2,3 | -68,4 | 10,6 |
| | Right Anterior Insula & Medial Frontal Lobe | 3149 | 43,2 | 15,3 | 8,2 |
| | Right Posterior Insula | 1277 | 48 | -24 | 13,7 |
| | Left Parieto-Occipital Cortex | 250 | -9,6 | -70,9 | 35 |
| | Right Parieto-Occipital Cortex | 245 | 12 | -68,7 | 35,6 |
| | Left Anterior Cingulate Cortex | 198 | -19,6 | 0 | 21,8 |
| | Left Somatomotor | 147 | -41,5 | -19,4 | 52,6 |
| | Right Hippocampus | 103 | 29,8 | -28,2 | -10,1 |
| | Right Anterior Cingulate Cortex | 69 | 18,7 | 18,6 | 16,5 |
| | Left Anterior Hippocampus | 61 | -26,1 | -16,5 | -19,7 |
| | Left Caudate | 57 | -13 | 13,1 | 13,9 |
| | Right Caudate | 52 | 12,5 | 18,7 | 9 |
| | Left Posterior Hippocampus | 51 | -24,9 | -37,6 | -2 |

| N3 SLOW WAVES | | | | | |
|---|---|---|---|---|---|
| **Signal Change** | **Region** | **Voxels** | **CM x** | **CM y** | **CM z** |
| Positive | Right Cerebellum | 607 | 26.3 | -45.6 | -26.5 |
| | Left Cerebellum | 276 | -24.4 | -44.1 | -26.8 |
| | Middle Cerebellum | 207 | -5.3 | -65.6 | -18.9 |
| Negative | Parieto-Occipital Cortex | 4456 | -0.8 | -64.6 | 18.9 |
| | Left Superior Temporal Lobe | 178 | -53.4 | -26.3 | 14.5 |
| | Left Orbitofrontal Cortex | 123 | -44.7 | 25.1 | -11.9 |
| | Bilateral Orbitofrontal Cortex | 67 | -4.1 | 49.6 | -10.2 |
| | Bilateral Somatomotor | 53 | -1.1 | -42.2 | 71.8 |

***Table S5. Brain areas showing significant BOLD-signal changes time-locked to the occurrence of slow waves in N1/N2 and N3 sleep, respectively (q<0.001 after FDR correction).*** *The table includes for each area the number of voxels and the coordinates of the center of mass (CM) in the standard MNI space (minimum cluster size = 50 voxels).*

**Figure S1**

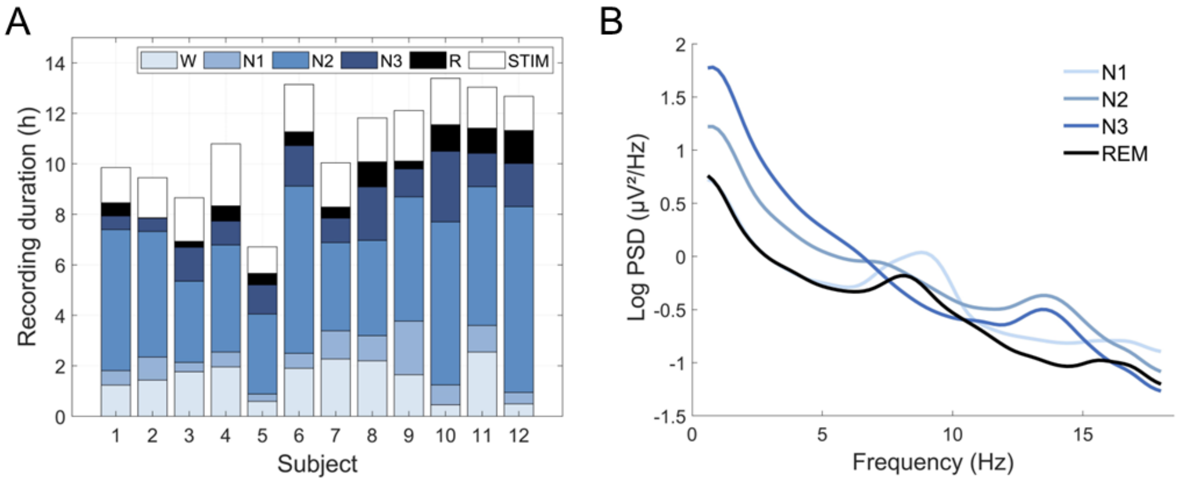


***Figure S1. Analysis of EEG recordings.*** *A. Amount of time spent in each sleep stage for each participant. Each column represents the sum of the two sessions for a different subject. STIM = artifactual segments, excluded from the analysis. B. Mean power spectral densities (PSD) obtained across all participants and recordings for N1, N2, N3 and REM sleep. PSD values were computed in two frontal (F3, F4) and two central (C3, C4) electrodes and then averaged.*

### Figure S2

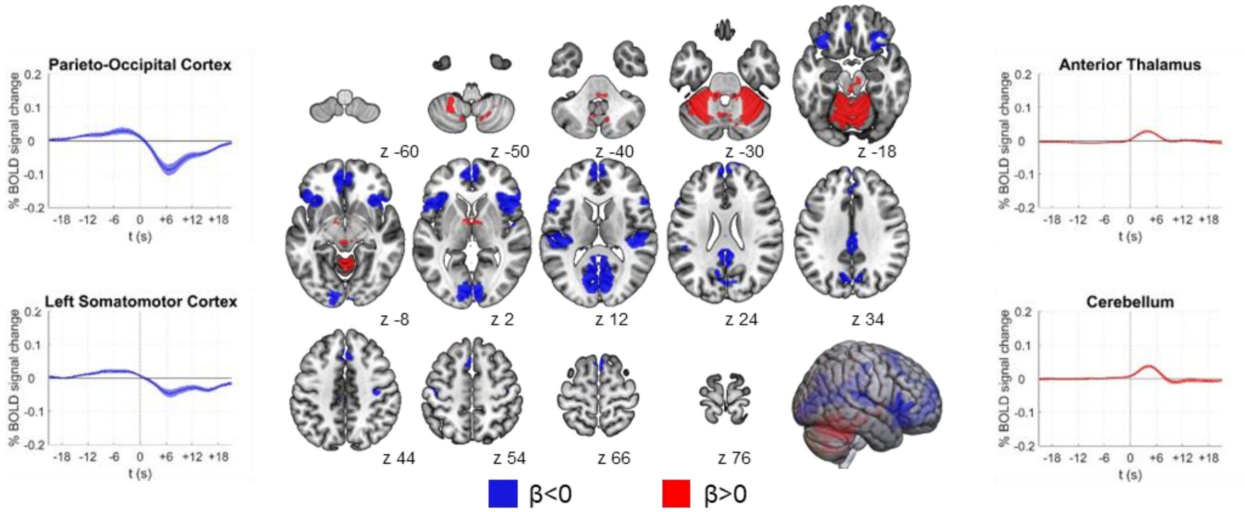


***Figure S2. Results of the regression analysis for slow waves.*** *Brain regions associated with a significant (q < 0.001) BOLD-signal increase are shown in red ($\beta > 0$), while regional BOLD-signal decreases are displayed in blue ($\beta < 0$). Brain images were generated using MRIcroGL (https://www.nitrc.org/projects/mricrogl/). Plots on the left and right show the mean BOLD-signals (up-sampled to the EEG sampling rate) time-locked to slow waves and the relative standard errors. These plots were obtained by averaging the signal of individual voxels within significant clusters identified through the regression analysis and cleaned out from sleep spindles signal contribution. t0: slow wave negative peak.*

### Figure S3

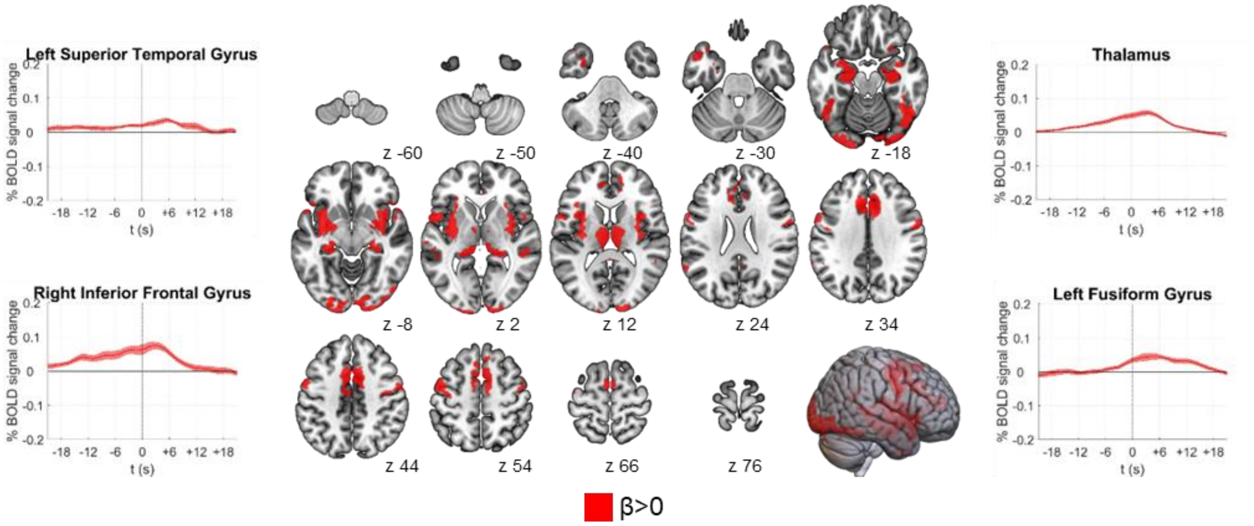


***Figure S3 - Results of the regression analysis for sleep spindles.*** *Brain regions associated with a significant ($q < 0.001$) BOLD-signal increase are shown in red ($\beta > 0$). Brain images were generated using MRIcroGL (https://www.nitrc.org/projects/mricrogl/). Plots on the left and right show the mean BOLD-signals (up-sampled to the EEG sampling rate) time-locked to sleep spindles and the relative standard errors. These plots were obtained by averaging the signal of individual voxels within significant clusters identified through the regression analysis and cleaned out from slow wave signal contribution. t0: sleep spindle onset.*

### Figure S4

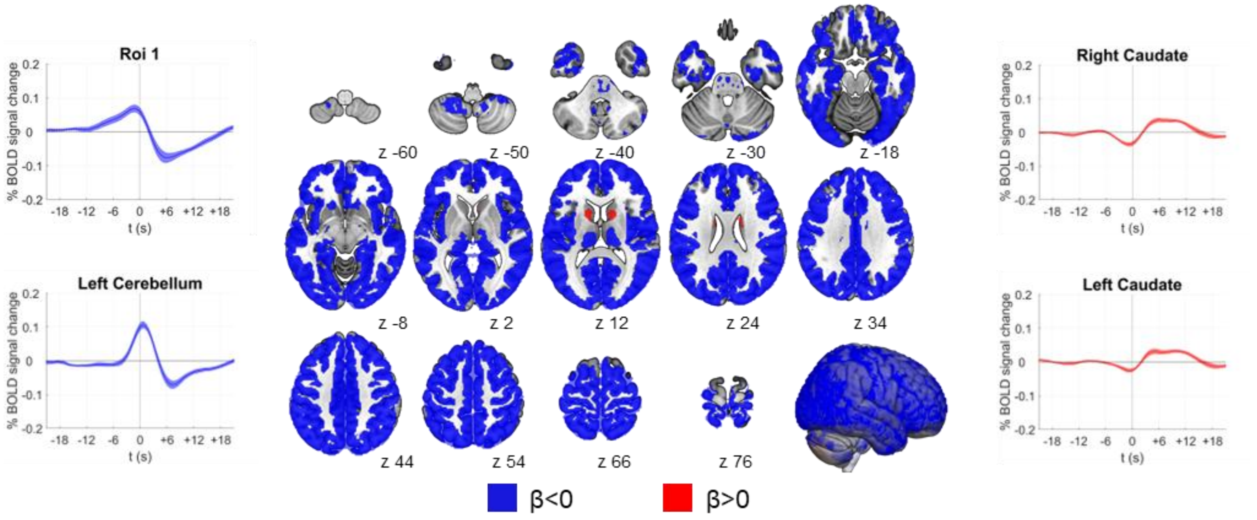


***Figure S4 - Results of the regression analysis for PWA-drops.*** *Brain regions associated with a significant (q < 0.001) BOLD-signal increase are shown in red (β > 0), while regional BOLD-signal decreases are displayed in blue (β < 0). Brain images were generated using MRIcroGL (https://www.nitrc.org/projects/mricrogl/). Plots on the left and right show the mean BOLD-signals (up-sampled to the EEG sampling rate) time-locked to PWA-drops and the relative standard errors. These plots were obtained by averaging the signal of individual voxels within significant clusters identified through the regression analysis and cleaned out from slow waves signal contribution. Roi 1 encompasses nearly all cortical areas, as well as specific subcortical structures, namely the thalamus and pons. t0: PWA-drop onset.*

**Figure S5**

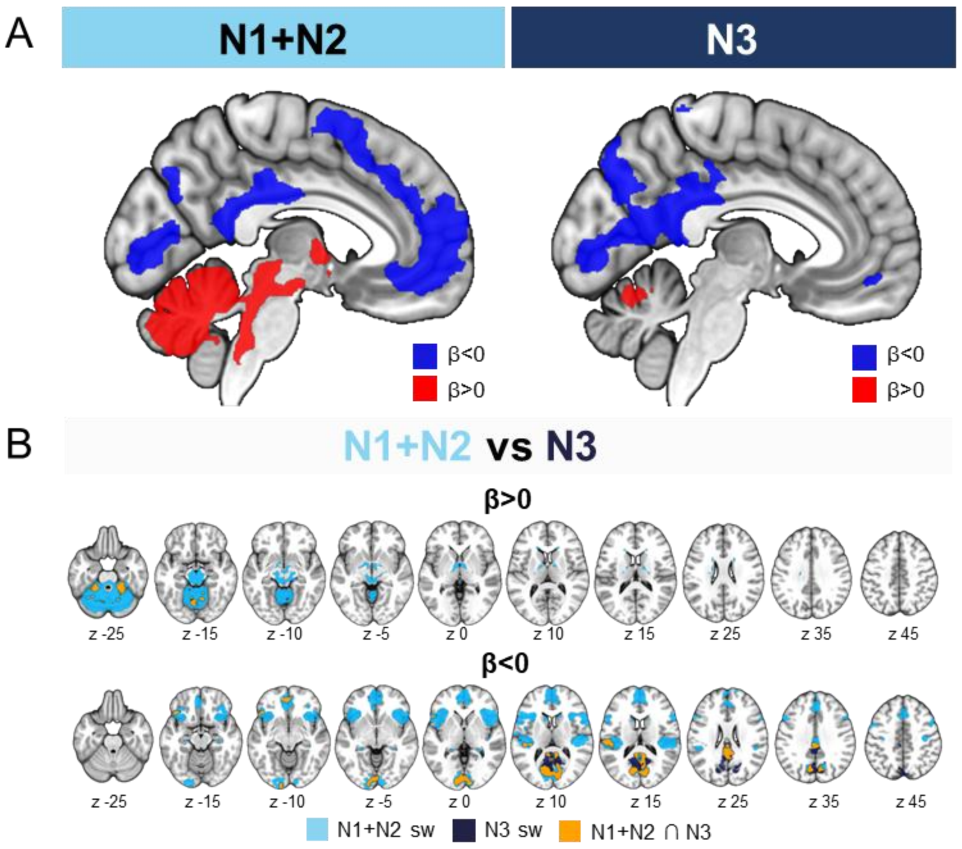


***Figure S5. Results of the regression analysis for N1+N2 and N3 slow waves.*** *A. Brain regions (q < 0.001) associated with a significant BOLD-signal increase are shown in red ($\beta > 0$), while regional BOLD-signal decreases are displayed in blue ($\beta < 0$). B. The axial brain slices illustrate the hemodynamic response patterns associated with light (N1+N2, in light blue) and deep (N3) NREM sleep slow waves (q<0.001, FDR corrected). Overlapping areas are shown in orange. Brain images were generated using MRIcroGL ([https://www.nitrc.org/projects/mricrogl/](https://www.nitrc.org/projects/mricrogl/) ).*

### Figure S6

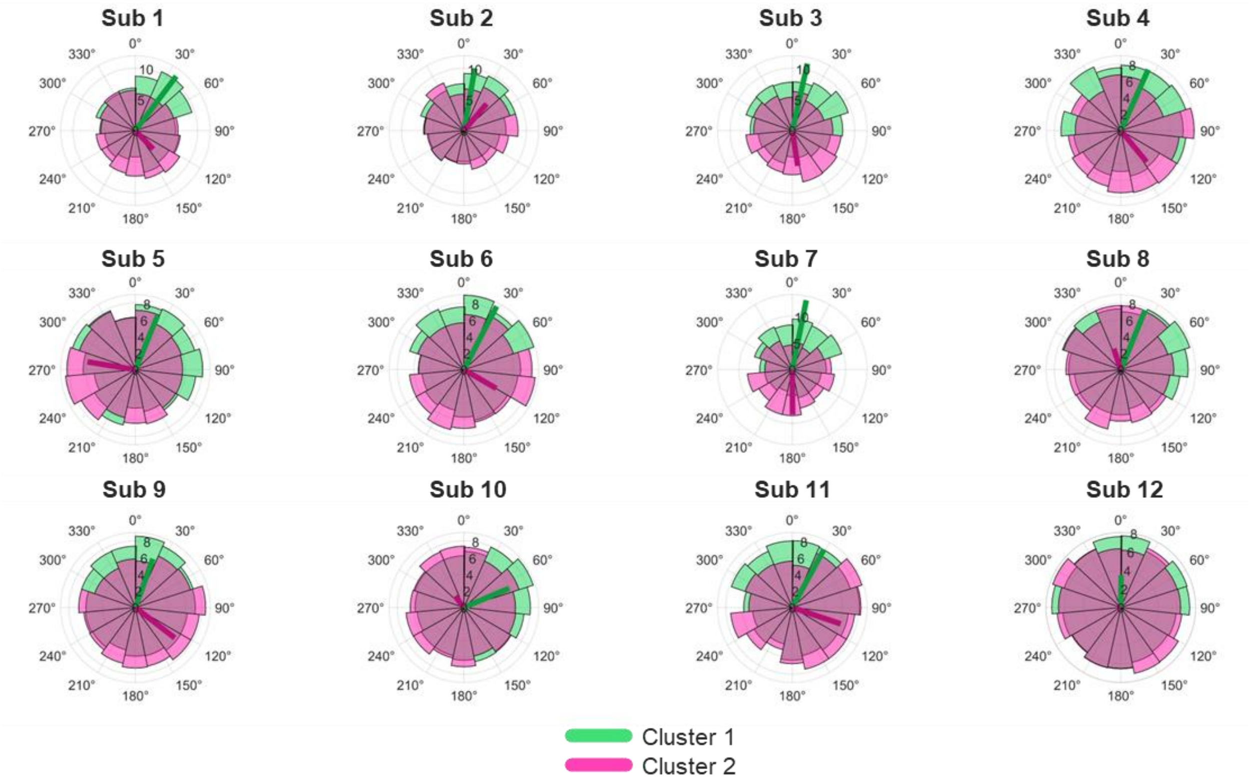


***Figure S6 - Polar plots illustrating the mean phase coupling at a single subject level between the onset of Cluster 1 (green) and Cluster 2 (pink) slow waves and the ISO phase, with 0° representing the ISO peak.*** *The plot includes the mean vector for each cluster. The full range of the infraslow phase was divided in 15 bins (24° each) for visualization purposes.*

**Figure S7**

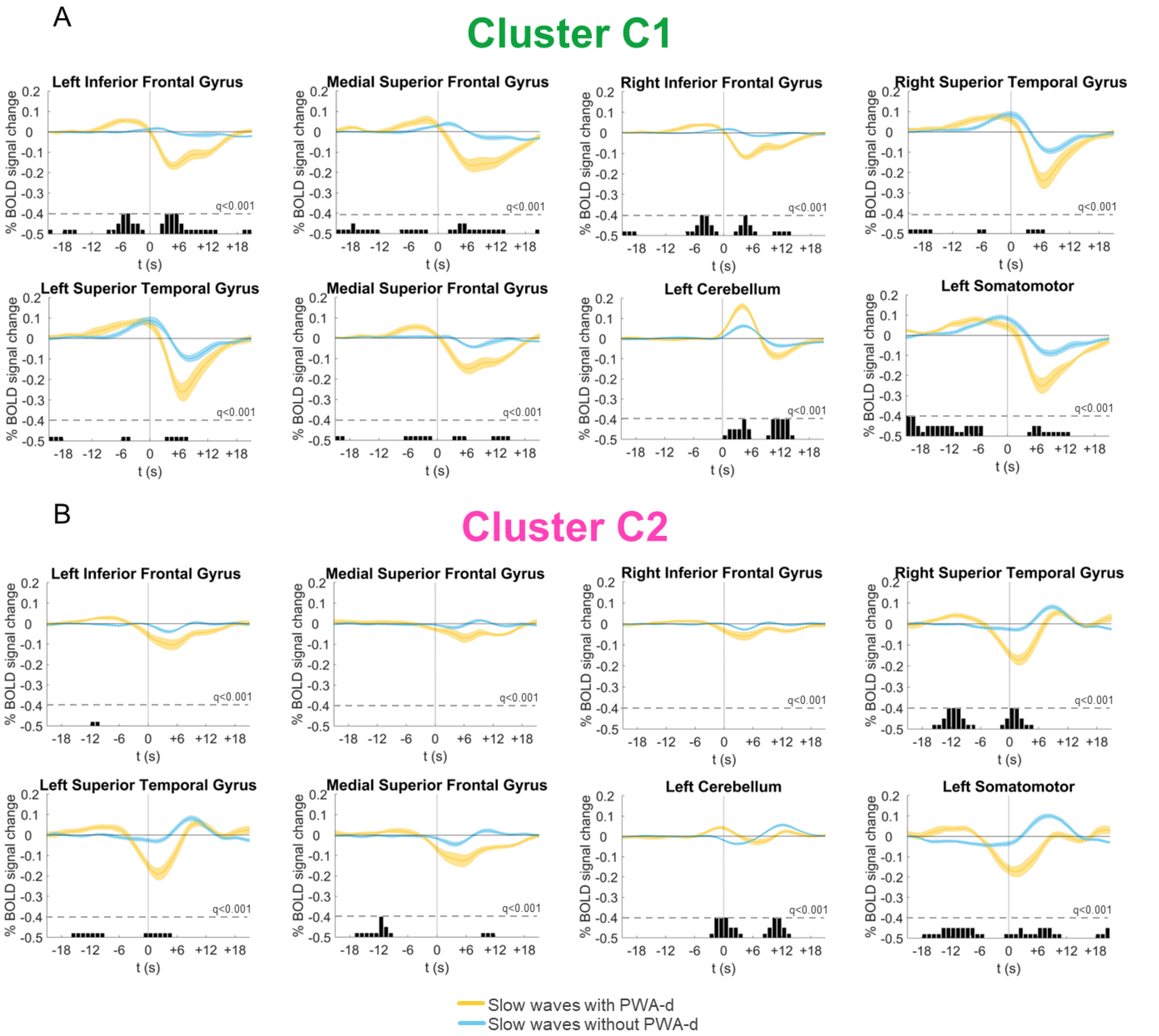


***Figure S7 - Mean BOLD signal changes time-locked to the onset of C1 (A) and C2 (B) slow waves are displayed, along with the corresponding standard errors****. These plots compare slow waves associated with a concurrent PWA drop (yellow) to those without a drop (light blue). Significant differences between the two conditions, calculated over 1-second bins (paired t-tests) and corrected for multiple comparisons using the FDR method, are indicated by black bars. The BOLD profiles were generated by averaging the signal across individual voxels within significant clusters identified through the main slow wave regression analysis.* ***$t_0$*** *: marks the negative peak of the slow wave.*
